# Genome Wide Association Studies on 7 Yield-related Traits of 183 Rice Varieties in Bangladesh

**DOI:** 10.1101/2020.11.22.393074

**Authors:** Nilanjan Roy, Acramul Haque Kabir, Nourin Zahan, Shahba Tasmiya Mouna, Sakshar Chakravarty, Atif Hasan Rahman, Md. Shamsuzzoha Bayzid

**Affiliations:** Department of Biomedical Engineering, Military Institute of Science and Technology, Bangladesh; Molecular, Cellular, and Developmental Biology, The University of Kansas, USA; Department of Biomedical Engineering, The University of Utah, USA; Department of Computer Science and Engineering, The University of California, Riverside, USA; Department of Computer Science and Engineering, Bangladesh University of Engineering and Technology

**Author notes:** Corresponding author: shams.

**Keywords:** Genome Wide Association Studies (GWAS), Rice, Yield-related traits, Single Nucleotide Polymorphism (SNP)

## Abstract

**Motivation:** Rice genetic diversity is regulated by multiple genes and is largely dependent on various environmental factors. Uncovering the genetic variations associated with the diversity in rice populations is the key to breed stable and high yielding rice varieties.

**Results:** We performed Genome Wide Association Studies (GWAS) on 7 rice yielding traits (grain length, grain width, grain weight, panicle length, leaf length, leaf width and leaf angle) based on a population of 183 rice landraces of Bangladesh. Our GWA studies reveal various chromosomal regions and candidate genes that are associated with different traits in Bangladeshi rice varieties. Noteworthy was the recurrent implication of chromosome 10 in all three grain shape related traits (grain length, grain width, and grain weight), indicating its pivotal role in shaping rice grain morphology. Our study also underscores the involvement of transposon gene families across these three traits. For leaf related traits, chromosome 10 was found to harbor regions that are significantly associated with leaf length and leaf width. The results of these association studies support previous findings as well as provide additional insights into the genetic diversity of rice.

**Conclusions:** This is the first known GWAS study on various yield-related traits in the varieties of *Oryza sativa* available in Bangladesh – the fourth largest rice-producing country. We believe this study will accelerate rice genetics research and breeding stable high-yielding rice in Bangladesh.

## 1 Introduction

Rice (*Oryza sativa* L.) is one of the most important food crops and feeds half the world’s population. Especially, this is the staple food of about 170 million people in Bangladesh, and this country has one of the highest per capita consumption of rice. Therefore, its food security largely depends on the good harvest of rice. Future increases in rice production, required to feed a continuously growing population of this country amidst various adverse climatic conditions due to climate change and limited arable land resources will rely primarily on genetic improvement of rice cultivars. Therefore, understanding the genetic basis of physiological and morphological variation in rice landraces in Bangladesh is critical for improving the quality and quantity of rice production. During the last few decades, great efforts in rice research have been made by Bangladesh Rice Research Institute (BRRI), in association with International Rice Research Institute (IRRI) to boost rice production. However, the current effort in increasing rice production in Bangladesh is mostly based on analyzing morphological characteristics and developing hybrids with *trial-and-error*. This traditional approach is not “scalable” to investigate the tremendous genetic and phenotypic variation of thousands of rice varieties available in Bangladesh. GWAS, considering the rice varieties in Bangladesh, may reveal important genotypephenotype associations which will direct the agricultural scientists towards a more informed research for breeding better rice varieties with desirable phenotypes suitable for the climate of Bangladesh.

Genome wide association studies (GWAS) have become a popular method to identify advantageous alleles and quantitative trait loci (QTL) associated with large-scale complex traits in rice population. Due to the growing awareness of the efficacy of GWAS in molecular dissection of traits and the abundance of genomic and phenotypic resources, many GWAS studies have been conducted over the past few years on various rice varieties across the world. Huang *et al.* (2010) [1] performed an association study on 14 rice agronomic traits across 373 indica rice varieties, and identified a total of 80 related sites. Huang *et al.* (2012) [2] identified a total of 32 heading date sites and 20 grain type sites across 950 rice varieties. Zhao *et al.* (2011) performed a GWAS on 34 traits across 413 rice varieties from 82 countries, and identified 234 associated sites [3]. Ya-fang *et al.* (2014) analyzed 315 rice varieties from the International Core Rice Germplasm Bank to perform a GWAS on five panicle traits, and a total of 36 candidate associated regions were detected [4]. Yang *et al.* (2014) performed GWAS on 15 traits, including 13 traditional agronomic traits and identified 141 associated loci [5]. Then they compared how these traits change along with the ecological environment. This led to the identification of valuable varieties and sub-groups with more favorable alleles. Biscarini *et al.* (2016) [6] conducted a genome-wide association analysis for grain morphology and root architecture for temperate rice accessions adapted to European pedo-climatic conditions, and a set of 391 rice accessions were GBS-genotyped leading to 57,000 polymorphic and informative SNPs. Among which 54% were in genic regions. A total of 42 significant genotype-phenotype associations were detected: 21 for plant morphology traits, 11 for grain quality traits, 10 for root architecture traits. The results helped them to dig into the narrow genetic pool of European temperate rice and to identify the most relevant genetic components contributing to a high yield of this germplasm. Zhang *et al.* (2019) [7] performed a GWAS with EMMAX for 12 agronomic traits using Ting’s core collection (7128 rice landraces from all over China and from some of the other main rice-cultivating countries collected by Ying Ting [8]). Yang *et al.* (2019) [9] detected SNP loci and determined related genes affecting the rice grain shape which lead to high-yielding breeding of rice. In that study, a total of 161 natural Indica rice varieties grown in southern China were used for a GWAS of grain shape-related traits. These traits include grain length (GL), grain width (GW), 1000-grain weight (TGW), and grain length/width (GLW). Ma *et al.* (2019) conducted a GWAS and a gene called OsSNB was identified controlling the grain size in rice [10]. Similarly, significant efforts have been made for association mapping with other yield related traits (e.g., panicle and leaf traits) [4, 5, 7, 9, 11].

In this study, we performed genome-wide association studies on 7 yield traits across 183 rice varieties in Bangladesh. We leverage the 3K Rice Genome Project (3K RGP) [12], where 3,000 rice genomes were re-sequenced and a resulting set of over 19 million SNPs has been characterized and made accessible [12–14]. While the previous GWAS studies provide fundamental resources regarding association mapping on various rice traits, none of them were especially targeted for Bangladeshi rice varieties. However, the grain yield-related rice traits are regulated by multiple genes, which are significantly influenced by the environment [1, 4, 15, 16]. As such, this study will further elucidate the impact of specific environmental conditions on the association between traits and genetic variations.

## 2 Materials and Methods

### 2.1 Rice materials

We leveraged the data from 3K RGP [12] (snp-seek.irri.org), which has sequenced a core selection of 3,000 rice accessions from 89 countries. We filtered a total of 183 rice varieties of Bangladesh. Detailed information on these 183 rice varieties are provided in Supplementary Material SM2. We considered seven yield-related phenotypes, namely grain length (GL), grain width (GW), grain weight (GWT), panicle length (PL), leaf length (LL), leaf width (LW) and leaf angle (LA). Out of the 183 rice varieties of Bangladesh in the 3K RGP, 168 had recorded phenotypic values for GL, GW, and GWT, while 158, 163, 160, and 84 rice varieties had values for PL, LL, LW, and LA, respectively [12]. Rice grain shapes are closely related to the yield and quality [9,17]. Leaf traits are among the major determinants of plant architecture, and are strongly associated to yield [18–21]. Panicle, being the top organ, is an important component in the canopy and is strongly correlated with spikelet yield [22]. Thus, investigating the genetic variations associated with these traits under specific conditions of Bangladesh would be fundamental to high-yield rice research in this country.

### 2.2 Characterization and genotypic data of rice germplasm

The characterization and breeding of various rice accession of Bangladeshi origin were performed over different years at IRRI, Los Baños, Philippines (14*^◦^* N, 121*^◦^* E, alt. 21 m) [23]. Genetic stocks for each of the 3,000 rice accessions were generated through one or more rounds of single-seed descent purification, conducted in field or screen-house settings [12]. Sowing occurred between May and July. Among the 3K RGP accessions, seeding date information was available for 29 Bangladeshi rice accessions, sown between 1989 – 2009 over different years, mainly between May and June (see Supplementary Figure S1). To capture diverse morphological and agronomic traits, characterization was conducted at three growth stages: vegetative, reproductive, and post-harvest. Leaf angle was measured during the vegetative stage, while panicle length, leaf length, and leaf width were measured in the reproductive stage. Grain trait-related information was collected in the post-harvest stage.

As part of 3K RGP, the 3,010 genomes were sequenced to an average depth of about 14x, ranging from approximately 4x to over 60x. Aligned with the Os-Nipponbare-Reference-IRGSP-1.0, the 3,010 genomes exhibited an average mapping coverage of 92% (range: 74.6% to 98.7%) [24]. SNP distribution varied widely among chromosomes, with chromosomes 4, 1, and 11 having the highest count and chromosomes 9, 10, and 5 having the lowest. Predominantly, SNPs were located in intergenic areas and introns, with only 18.24% in exons, almost 40% of which were synonymous. A total of over 29 million SNPs were discovered, with over 27 million being bi-allelic and exhibiting strong concordance (*>* 96%). Post-filtering, a core set of about 17 million SNPs was obtained covering nearly 99.9% of all SNPs with a minor allele frequency exceeding 0.25%. Notably, 56% of non-transposable element (NTE) genes and 91% of transposable element (TE)-related genes contained high-effect SNPs [25].

### 2.3 Data analysis

Various statistical analyses of the yield-related traits were performed using R [26]. We used raw phenotypic values for these statistical analyses. We computed Broad-sense heritability, *H*^2^. The Shannon diversity index for traits was computed with the vegan package in R [27].

To address the impact of covariates like environment, subpopulation, and seeding date on trait phenotypic values, we applied the Best Linear Unbiased Prediction (BLUP) model using the lmer4 package in R [28]. Given the multi-environmental nature of the data used in our GWA studies, we attempted to account for potential confounding factors by including subpopulation and seeding date as covariates in the model to adjust the phenotypic values of Bangladeshi rice varieties. Our model treated rice varieties as fixed effects, while subpopulation and seeding date were considered random effects, thereby providing adjusted phenotypic values for our analysis. We used the following model where (1*|s*) denotes random intercepts for each unique level of the factor *s*. *Y* and *X* indicate the adjusted phenotypic BLUP value and raw phenotypic value of the rice varieties respectively.

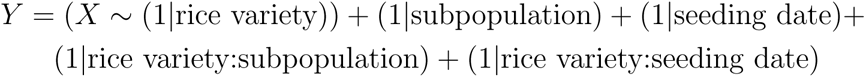

To understand the genetic diversity within Bangladeshi rice varieties and the diversity compared to the 3K RGP panel, we calculated minor allele frequency (MAF), nucleotide diversity *π* and Tajima’s D in the 3K RG 404K CoreSNP dataset (snp-seek.irri.org). In the 3K RGP dataset, 404K core SNPs were filtered using a two-step linkage disequilibrium (LD) to limit the number of markers with *MAF >* 0.01 and a call rate of 0.8. We used PLINK (freq) to calculate the MAF [29]. Nucleotide diversity *π*, and Tajima’s D were calculated with VCFtools [30]. We used sliding windows of 1000kb and 100kb for calculating the nucleotide diversity *π*, and Tajima’s D, respectively, within the Bangladeshi rice varieties and compared to the 3K RGP panel.

For GWAS, the 3K RG 29mio biallelic SNPs dataset was processed using PLINK. Filtering SNPs (missing rate *<* 50% and MAF *>* 0.05) to reduce the false-positive rate resulted in 39,40,165 SNPs. A Hardy–Weinberg equilibrium filter (*p <* 10*^−^*^6^) was applied to exclude markers that deviate from Hardy–Weinberg equilibrium and to account for genotyping error, and exclusion of samples with kinship coefficient *>* 0.177 was applied to account for the popultion stratification effects (i.e., to remove first-degree relationships of the rice varieties). After excluding first-degree relationships, sample counts were 151 (GL, GW), 154 (GWT), 139 (LL), 143 (LW), 74 (LA), and 129 (PL).

To address linkage disequilibrium (LD), we pruned variants using PLINK. LD was computed within 1500-SNP windows, and LD *>* 0.6 (GL, GW, LW, LA) or *>* 0.7 (GWT, PL, LL) resulted in SNP exclusion. Principal Component Analyses (PCA) were then performed to explore population structure. This process yielded 420,519 SNPs for GL and GW, 514,640 for GWT, 258,323 for LL, 420,375 for LW, 407,435 for LA, and 178,543 for PL.

The quantitative association test function of PLINK (–*glm*) was used to get the *P* - values of significant SNPs. To correct for population structure, normalized PC1 and PC2 were included as covariates in the association tests. The genomic inflation factor (*λ*) was computed using the *glm* function for each GWAS feature, which quantifies the amount of the bulk inflation as well as the excess false positive rate, with values ranging from 1 to 1.10, indicating no evidence of inflation [31]. A significant threshold of *−* log_10_ *P* was set, where 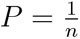, with *n* being the total number of markers used [7, 9].

Considering that GWAS hits are often not causative, but rather SNPs in LD with them, we performed genome-wide LD analysis on datasets that are not LD-pruned prior to association tests. LD decay in cultivated rice is typically between 100-200 kb [32]. For GL, we extracted SNPs within *±*100 kb of GWAS hits; for other traits, *±*50 kb. PLINK was used to extract SNPs around GWAS hits, and Haploview for genome-wide LD analysis [33]. LD plots were created using the R package LDheatmap [34]. Annotation analysis utilized SnpEff results of 3K RGP 29mio biallelic SNPs from 3K RGP [12] (snp-seek.irri.org), while candidate gene information was sourced from the Michigan State University (MSU) Rice Genome Annotation Project Database [35]. Data visualization was done using the Tidyverse package in R [36].

## 3 Results and Discussion

### 3.1 Phenotypic diversity of the traits

Statistical analyses of the phenotypic diversity in the yield-related traits are shown in Table 1. The minimum and maximum GL in this population are 4.7 and 10.2, respectively, having a mean GL of 8.28 with 9.48% coefficient of variation. The CV values of various grain shape related traits range from 9.48% *∼*16.4%, and those of leaf and panicle traits range from 11.67% *∼*57.7%. These analyses suggest that this group of rice varieties represents significant variations in grain shape and other yield-related traits.

**Table 1:** Statistical analyses of the yield-related traits. We show the minimum (Min), maximum (Max), mean, standard deviation (SD), coefficient of variation (CV), heritability and Shannon diversity index for each of the 7 traits.

| Trait | Min | Max | Mean | SD | CV (%) | Heritability | Shannon<br>diversity<br>index |
| --- | --- | --- | --- | --- | --- | --- | --- |
| GL (mm) | 4.7 | 10.2 | 8.28 | 0.78 | 9.48 | 0.47 | 1.298754 |
| GW (mm) | 2.1 | 3.9 | 3.05 | 0.33 | 10.8 | 0.44 | 1.298754 |
| GWT (gm) | 1.2 | 3.5 | 2.4 | 0.4 | 16.4 | 0.52 | 1.298754 |
| LL (cm) | 2 | 5 | 3.26 | 0.63 | 19.41 | 0.68 | 1.3 |
| LW (cm) | 0.8 | 2.3 | 1.29 | 0.24 | 19.26 | 0.73 | 1.32 |
| LA (rd) | 1 | 9 | 3.39 | 1.96 | 57.7 | 0.49 | 1.45 |
| PL (cm) | 18 | 32 | 24.89 | 2.90 | 11.67 | 0.42 | 1.3 |

The distribution of these traits as well as the correlations between them are depicted in Fig. 1. It suggests that these traits are normally distributed, and GL, GW and GWT are positively correlated with each other, especially GW and GWT are strongly correlated with each other. To account for the impact of various factors such as environment, subpopulation, and seeding date, we adjusted the data using BLUP. The distribution of the BLUP-adjusted data is presented in supplementary material. The linear relationship between the adjusted phenotypic values and the raw phenotypic values (presented in Supplementary Figures S11 and S12) suggests that we can utilize the adjusted phenotypic values for our GWAS analysis.

**Figure 1:**
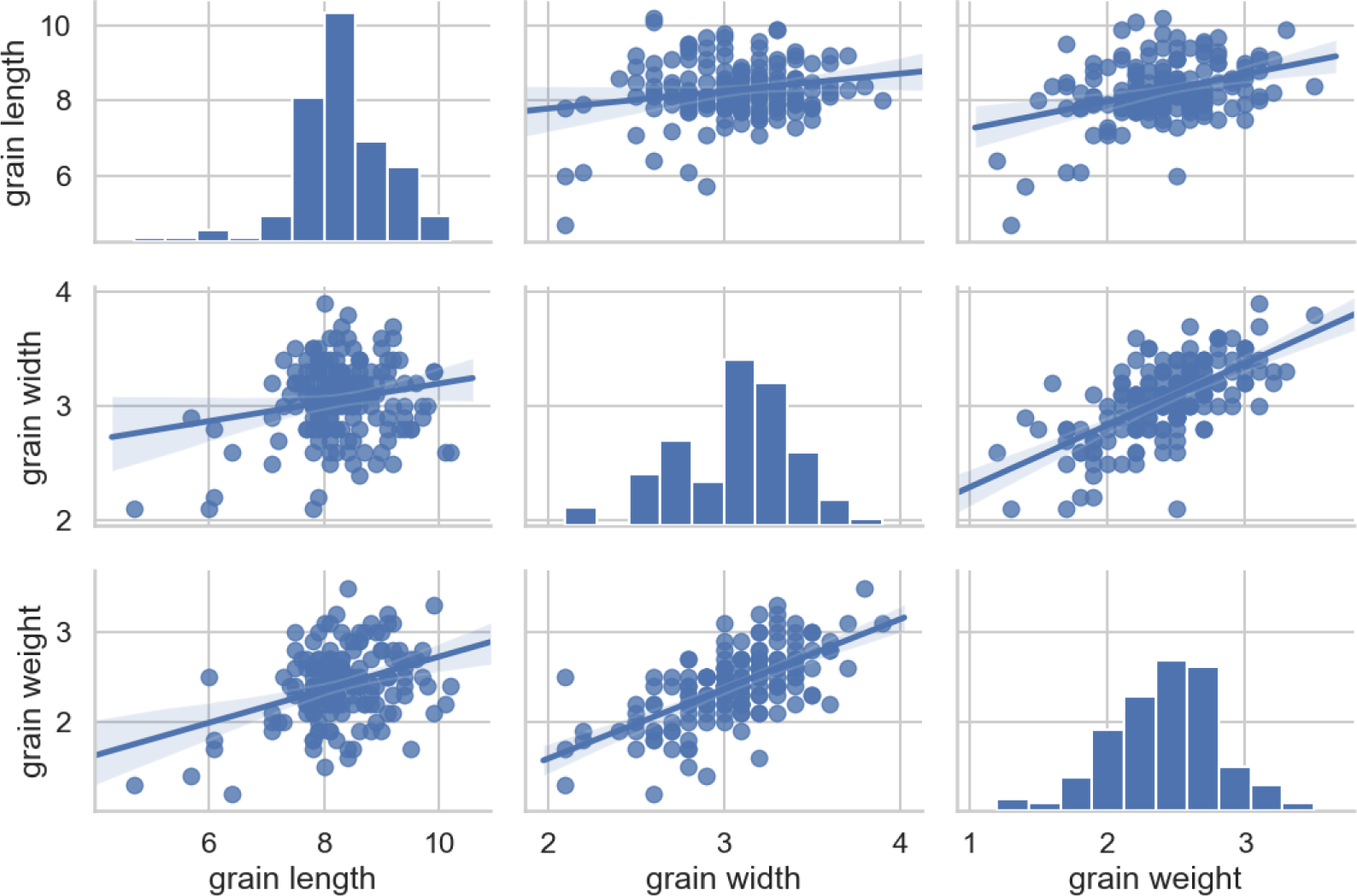
Statistical analyses of the grain shape related traits. We show the distribution and correlation scatter matrix of these three traits. GL and GW are shown in millimetre (mm) and GWT is shown in gram (gm). The blue lines in the scatter plots indicate the correlation trends.

We also calculated broad sense heritability, *H*^2^ and Shannon diversity index of the traits analyzed in this study. The values are shown in Table 1. The heritability values indicate that these traits are highly heritable and genetic influence is moderately high. The heritability of these traits, reported in previous studies [37, 38], appears to be higher than what we observed in our study of Bangladeshi rice varieties. This suggests a significant influence of environmental and other factors on these traits within Bangladeshi rice varieties. The Shannon diversity indices indicate high diversity in the samples analyzed in this study.

### 3.2 Genetic diversity analysis

In Figure 2, we present MAF distributions, comparing the MAF distributions of 3K RGP lines with those of Bangladeshi lines, and among the subpopulations of Bangladeshi varieties. Notably, 3K RGP lines show a higher density of low-frequency allele variation compared to Bangladeshi lines (Figure 2(a)). Moreover, 3K RGP lines exhibit a higher distribution of variants with a frequency of 0.5. Figure 2(b) highlights ARO as having the highest prevalence of low-frequency allele variation among Bangladeshi subpopulations, followed by AUS. Both ADMIX and IND subpopulations show similar, albeit lower, densities of low-frequency allele variation compared to ARO and AUS. Notably, ADMIX and ARO subpopulations exhibit distinct peaks in the minor allele distribution plot, suggesting unique genetic variants at specific loci, which is not observed in AUS and IND subpopulations.

**Figure 2:**
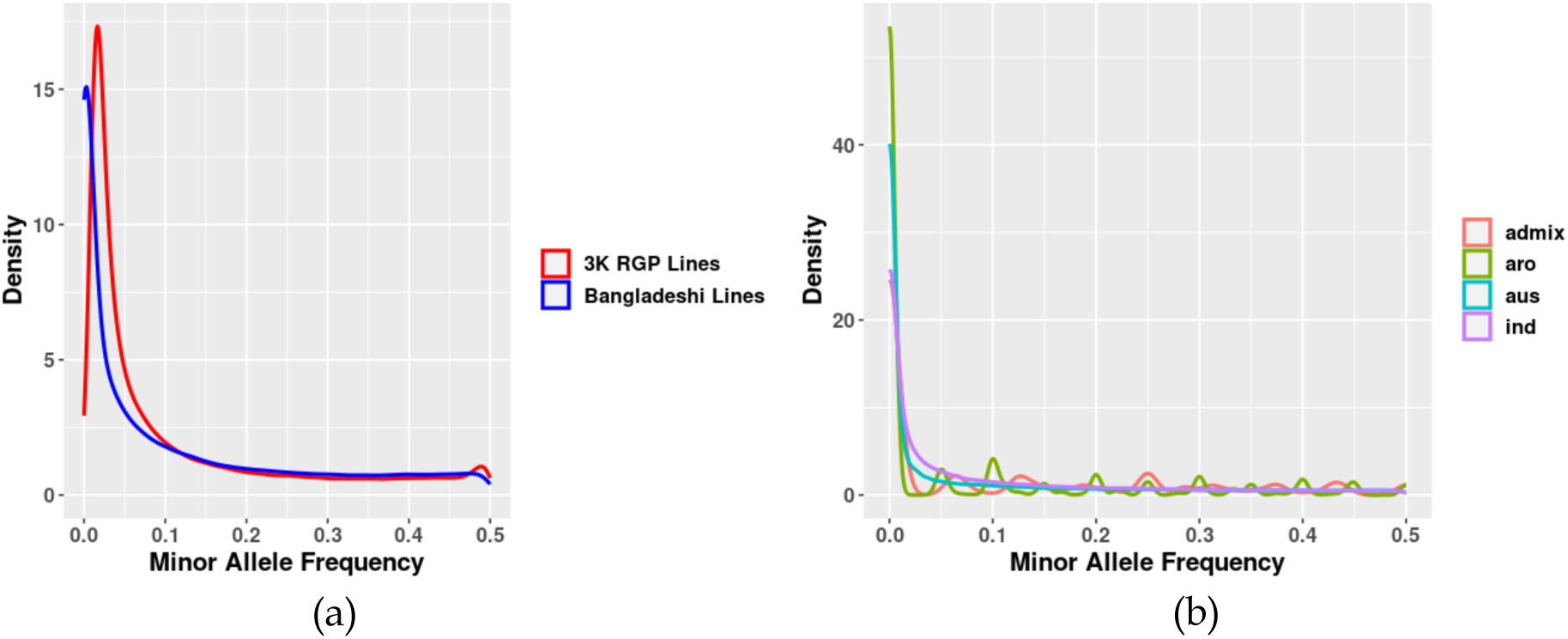
Density distribution of minor allele frequencies in rice populations. (a) The distribution of minor allele frequencies of the 3K RGP lines (excluding Bangladeshi varieties) with all the Bangladeshi lines. (b) The distribution of minor allele frequencies among the Bangladeshi lines.

Figure 3 shows a comparative analysis of nucleotide diversity (*π*) between the 3K Rice Genome Project and Bangladeshi rice lines. Nucleotide diversity is computed along the length of each chromosome, measured in megabases (Mb). While the overall chromosomewide nucleotide diversity trends are akin between the 3K RGP and Bangladeshi lines, Chr 10 exhibits notably less diversity in the Bangladeshi lines in contrast to the 3K RGP lines. This may suggest that Chr 10 is more conserved in the Bangladeshi lines. Examining specific regions, such as those near 15, 25, 40 Mb in Chr 1; 5 Mb in Chr 2; 5, 20, 30 Mb in Chr 3; 2, 15, 20 Mb in Chr 4; 15 Mb in Chr 5; 1, 10, 24 Mb in Chr 6; 21, 28 Mb in Chr 7; 10, 18 Mb in Chr 8; 12 Mb in Chr 9; 22 Mb in Chr 11; 17 Mb in Chr 12, reveals higher nucleotide diversity in 3K RGP lines compared to Bangladeshi lines. This suggests a lower level of variation in these regions within the Bangladeshi lines. Conversely, regions near 25 Mb in Chr 2; 15, 35 Mb in Chr 3; 22, 32 Mb in Chr 4; 1 Mb in Chr 5; 18, 30 Mb in Chr 6; 15, 22 Mb in Chr 7; 8, 30 Mb in Chr 8; 7 Mb in Chr 9; 20 Mb in Chr 20 exhibit higher nucleotide diversity in Bangladeshi lines, implying possible diversity in Bangladeshi rice varieties in these regions.

**Figure 3:**
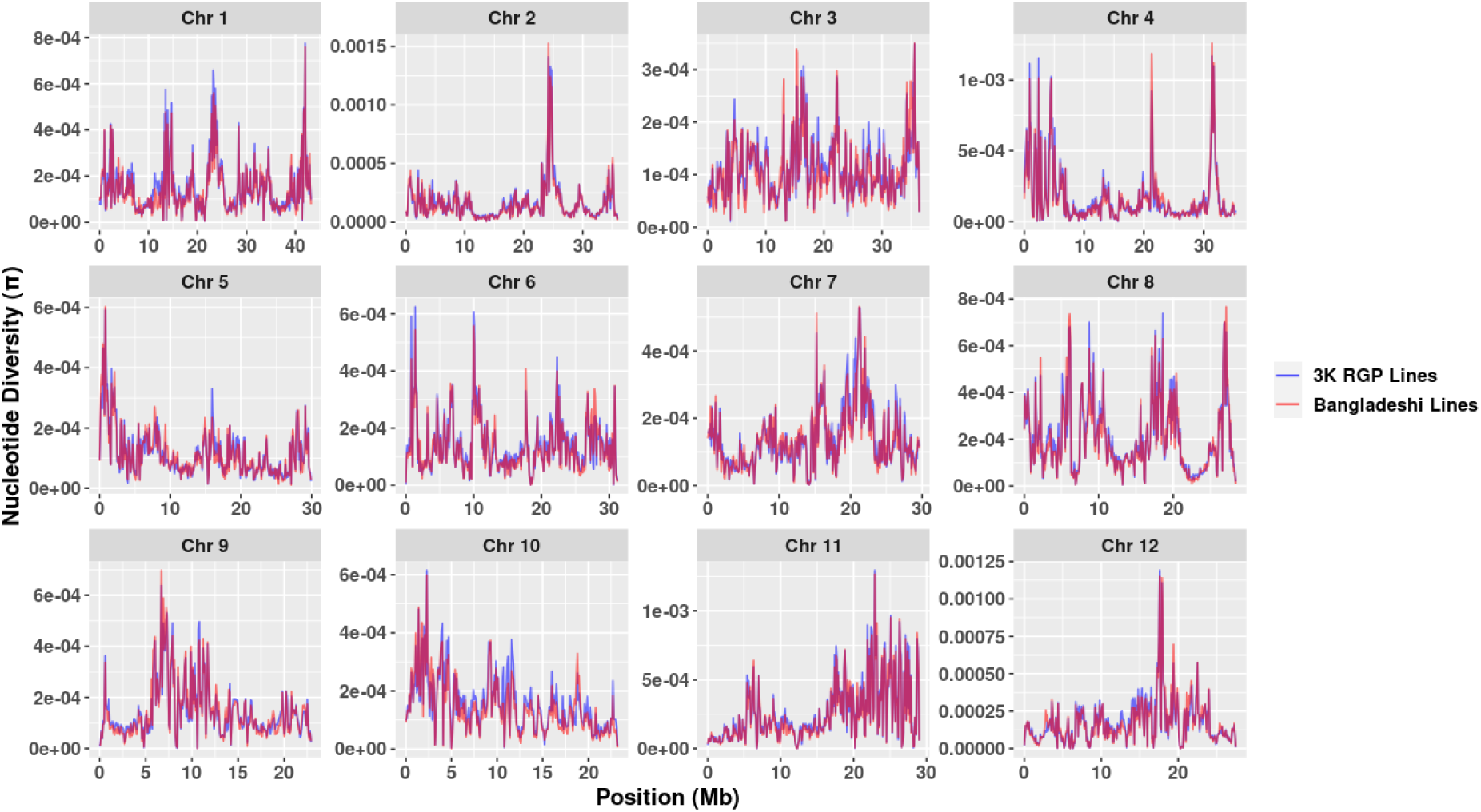
Nucleotide diversity (*π*) across rice chromosomes for 3K RGP and Bangladeshi lines.

Figure S2 in the supplementary material illustrates the nucleotide diversity across chromosomes in different subpopulations of Bangladeshi rice. Chromosomes 2, 4, 11, and 12 show similar nucleotide diversity patterns among subpopulations, indicating uniform allelic variations within these chromosomes. However, specific regions, such as those near 1-10, 20, 30 Mb in Chr 2; 7, 12 Mb in Chr 4; 8, 25 Mb in Chr 11; 5-10, 22 Mb in Chr 12, exhibit variations in nucleotide diversity among subpopulations. In contrast, other chromosomes display varying nucleotide diversity patterns. Specifically, regions near 30 Mb in Chr 1; 15-30 Mb in Chr 3; 0-10 Mb in Chr 5; 8, 12, 22 Mb in Chr 6; 20 Mb in Chr 7; 0-5, 8-15, 18, 28 Mb in Chr 8; 10, 12-30 Mb in Chr 9; 3-18 Mb in Chr 10 exhibit distinct nucleotide diversity among subpopulations.

Additionally, we computed genome-wide Tajima’s *D* to identify and compare selection signatures between 3K RGP and Bangladeshi lines (Figure S3 in the supplementary material). A notable divergence in selection patterns was observed in Chr 5, specifically in the 8-14 Mb region. Here, 3K RGP lines showed an excess of rare alleles (Tajima’s *D <* 0), while Bangladeshi lines demonstrated a scarcity of rare alleles (Tajima’s *D >* 0). Further investigation within Bangladeshi lines revealed distinctive selection patterns among subpopulations (Figure S3(b)), notably in regions near 8-15 Mb in chromosome 2 and 10-15 Mb in chromosome 8.

### 3.3 Population structure

Principal component analysis (PCA) was performed based on 39,40,165 SNPs in 183 rice varieties. The first and second PCs captured 35% and 23% of the total variation respectively, indicating its highly structured nature. PCA plot of the first two principal components is shown in Fig. 4 which suggests three major subgroups: indica (IND), aus (AUS), and aromatic (ARO). The principal components representing the IND samples are located in the lower left, the ARO samples are located in the upper right, and the AUS sample are located in the lower right part. There are a few ad-mixed (intermediate type) varieties as well. We also show the genome stratification based on SNP markers using multidimensional scaling (MDS) plot (see supplementary Fig. S4).

**Figure 4:**
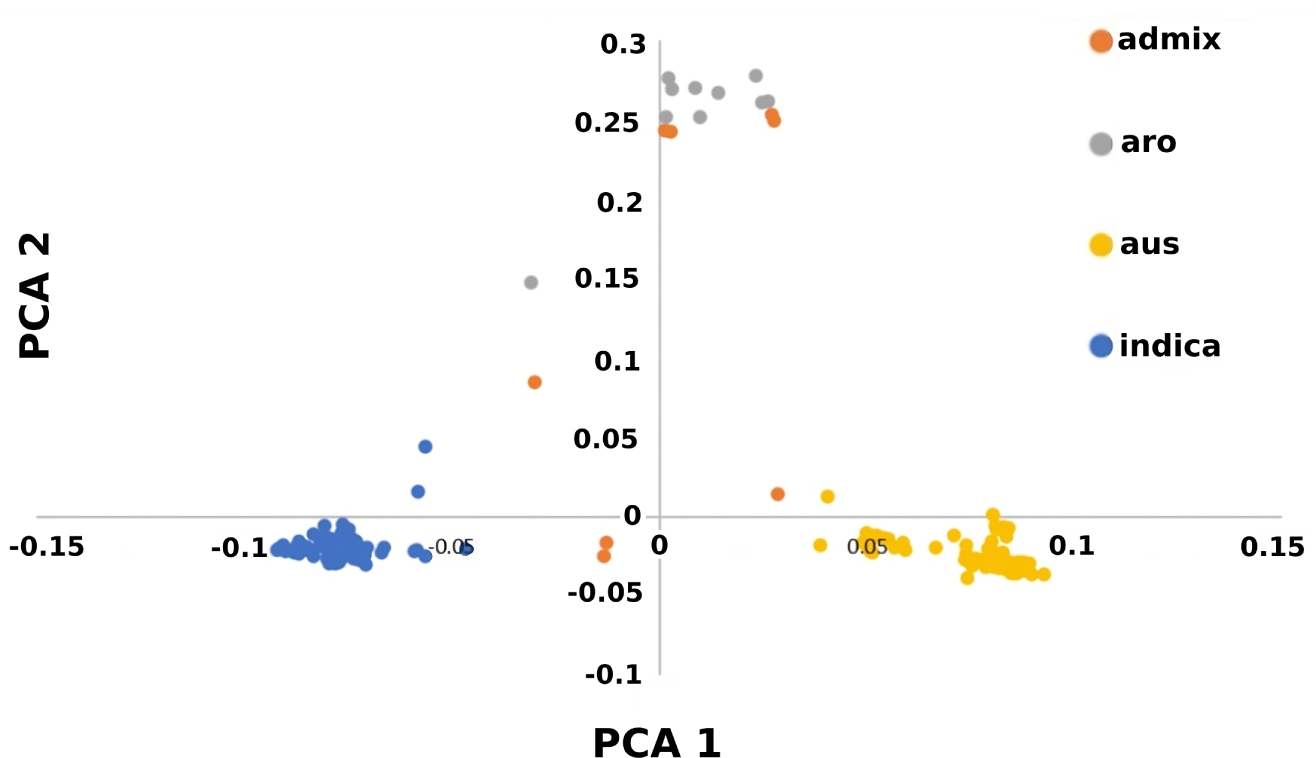
Population structure analysis using PCA. We show the PCA plot of the first two principal components of 183 rice varieties.

Since population stratification is observed in the samples, we adjust for population stratification while testing for associations between the SNPs and the phenotypes. We observed that the first two principal components capture 58% of the variation as well as the three main subgroups of rice varieties in our samples. So, we use the first two axes of variation as covariates in the *P* -value calculation.

### 3.4 Relative kinship among the rice varieties

The kinship matrix was calculated using a centered IBS (identity by state) method [39] in TASSEL [40]. Notably, most of the kinship estimates were zero (or close to zero) and only a few kinship values were above 0.5. It suggests that these varieties are reasonably unrelated and that there is a lower possibility of any spurious relationships confounding GWAS outcomes [7]. Figure. 5 shows the kinship matrix as well as histograms of the realized relationship coefficients. The distributions of off-diagonal and diagonal elements (Figs. 5(b)-(c)), indicate a highly structured rice population which was also supported by our PCA analyses reported in Sec. 3.3.

**Figure 5:**
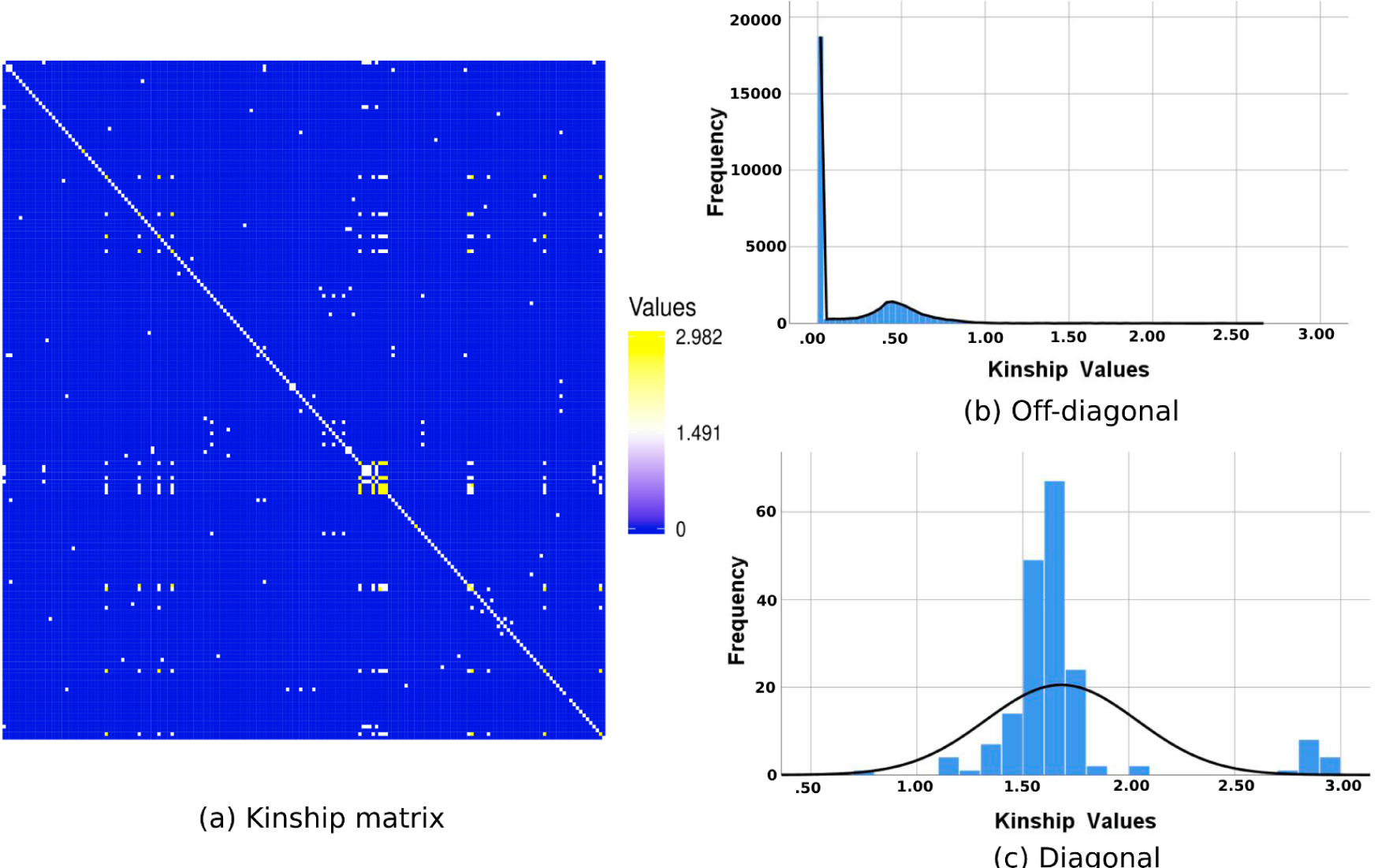
Relative kinship among 183 rice varieties. (a) A heatmap representing the kinship coefficients. (b)-(c) Histograms of the realized relationship coefficients, where we show the frequencies of the off-diagonal and diagonal elements in the kinship matrix respectively.

### 3.5 GWAS on grain shape related traits

A total of 168 rice varieties (out of 183 Bangladeshi varieties collected from the 3K RGP) had GL, GW and GWT values associated with them, and we used these 168 varieties for our association studies. To delineate genetic determinants of various traits in rice, we identified significant SNPs from Manhattan plots. The genomic inflation factor was calculated to test the accuracy of the GWAS models and to mitigate false positives. In addition, Q-Q (quantile-quantile) plots were used to check whether the p-values are inflated. As GWAS hits may not necessarily be within the causal genes, we executed a genome-wide linkage disequilibrium (LD) analysis for the candidate peak regions to characterize regions harboring putative candidate genes [41]. LD analyses were done with the datasets that are not LD pruned while GWAS analyses were done with LD pruned data. These candidate peak regions (SNPs that are in LD with GWAS hits) are provided as supplementary data files (SM4-SM10). We have concentrated our LD analyses on SNPs that are correlated with GWAS hits and possess functional annotations affecting the traits we’re interested in.

#### 3.5.1 GWAS for grain length

The Manhattan plot shown in Figure 6 reveals three noteworthy SNPs harbored in chromosomes 3, 7, and 10, corresponding to genes LOC Os03g38850, LOC Os07g11640, and LOC Os10g22620, respectively (see Table 2). Notably, these three genes encode retrotransposon proteins, a class of transposable elements known for their ability to mobilize within and across genomes, thereby contributing to genomic diversity and size variation. In the realm of plant biology, Long Terminal Repeat (LTR) retrotransposons, a subtype of retrotransposons, have been recognized as significant contributors to genome evolution [42]. LTR retrotransposons are not only important for gene regulation but also play a crucial role in plant heat response [43].

**Figure 6:**
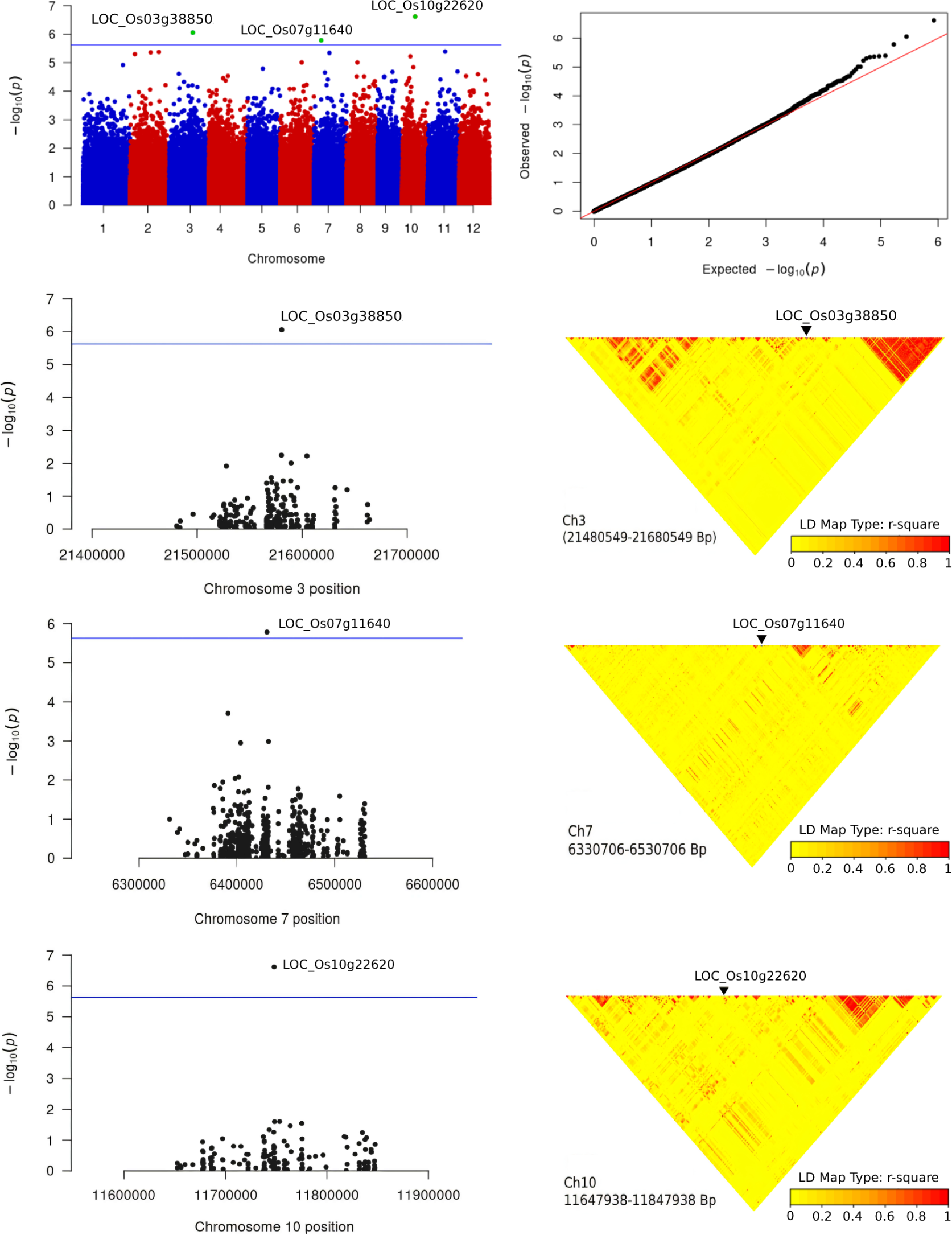
GWAS on grain length using 168 rice varieties. We show the Manhattan plots, Q-Q plots, local Manhattan plots, LD heatmap plots. The blue lines indicate the genome-wide significant threshold. For QQ plots, the horizontal axis shows the *−* log10-transformed expected *P* -values, and the vertical axis indicates *−* log10-transformed observed *P* -values. We color the LD heatmaps with a color gradient which varies continuously from yellow to red with increasing *r*^2^ values.

**Table 2:**
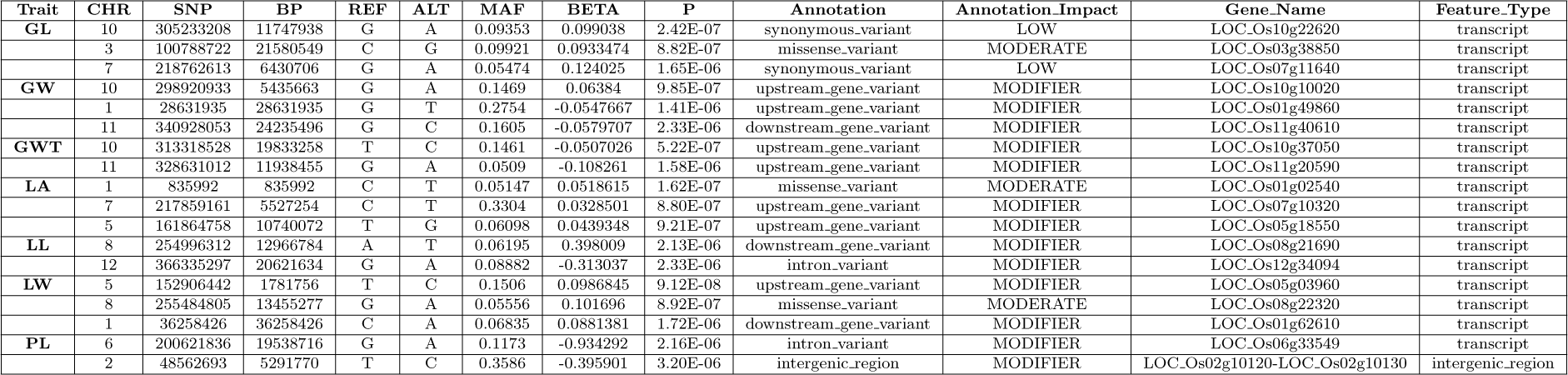
Genome-wide lead SNPs significantly associated with various traits.

While the specific involvement of retrotransposon proteins in grain length remains implicit in the previous studies, the broader literature underscores their impact on rice grain-related traits. A noteworthy example is the association of LTR retrotransposons with grain width, leading to the development of extra-large grains and a substantial increase in rice yield [44]. Thus, their potential implications for key agronomic traits, including grain length, cannot be overlooked.

In our LD analysis aimed at exploring additional alleles proximal to the significant genes, an array of genes associated with grain length regulation was identified. Notably, LOC Os03g38710 (harbored in chromosome 3) was identified as encoding a serine-threonine protein kinase, suggesting potential involvement in signaling pathways pivotal to the regulation of grain length. Additionally, a cluster of genes – LOC Os03g38830, LOC Os03g38840, LOC Os03g38850, LOC Os03g38880, and LOC Os03g38890 – was found to encode retrotransposon proteins, suggesting significant impact on grain lengths other grain traits in rice [45]. Another noteworthy gene is LOC Os03g38930, which encodes a protein featuring a signal peptide. Previous studies suggest that signal peptides such as epidermal Pattering Factor-Like2 (EPFL2) regulate grain number, grain length, and awn development in rice [46, 47].

Our analysis also revealed a series of genes harbored in chromosome 10 – LOC Os10g22550, LOC Os10g22540, LOC Os10g22620, LOC Os10g22640, LOC Os10g22650, LOC Os10g22760, and LOC Os10g22800 – that are known to encode retrotransposon proteins.

The gene LOC Os10g22560 encodes a peptide transporter PTR2n, which is involved in the transport of nitrogen-containing substrates [48]. Although not directly linked to grain length, nitrogen transport and metabolism can indirectly influence grain development and size. In rice, the PTR gene family to which LOC Os10g22560 belongs is involved in nitrate uptake and transport [49]. The expression profiles of 96 PTR genes in rice have been analyzed, showing their potential role in nitrogen use efficiency (NUE) and nitrogen metabolism pathways [49]. Therefore, while not directly linked to grain length, the gene LOC Os10g22560 and the PTR gene family are important for nitrogen transport and metabolism, which can influence grain development and size.

Further exploration of chromosome 7 revealed LOC Os07g11500, LOC Os07g11640, LOC Os07g11650, LOC Os07g11670, LOC Os07g11700, LOC Os07g11720, LOC Os07g11790, and LOC Os07g11800 were found to encode retrotransposon proteins, aligning with their potential impact on grain length and other grain traits in rice [50].

Our GWAS study supports previous findings as well as reveal additional chromosomal regions associated with GL. Yang *et al.* (2019) [9] identified eight SNP loci in four chromosomes (3, 5, 6, and 7) to be in close association with grain length, and – among these four chromosomes – our study identified chromosomes 3, 7 to harbor two significant loci. Similarly, Ya-fang *et al.* (2015) identified 10 SNP loci in choromosomes 3, 5, 6, 7, 8, 10, and 12, which were significantly associated with GL [4]. A study on basmati rice of Indian origin (Singh *et al.* (2012) [51]) also revealed chromosomal region 7 to be associated with GL, and an SSR (simple sequence repeat) analysis in Pakistan (Aslam *et al.* (2014) [52]) identified chromosomal regions in 3 and 7 to be in close association with grain length.

#### 3.5.2 GWAS on grain width (GW)

GWAS on GW revealed a triad of significant SNPs on chromosomes 1, 10, and 11, aligning with genes LOC Os01g49860, LOC Os10g10020, and LOC Os11g40610, respectively, as depicted in the Manhattan plots of Fig. 7(a). The gene LOC Os11g40610 in chromosome 11, encoding an early flowering protein, emerged as the most important candidate for grain width trait as early-morning flowering in rice influences grain yield [53]. When we looked at the other two GWAS hits in chromosomes 1 and 10 in the Rice Genome Annotation project (http://rice.uga.edu/), we observed that these are conserved proteins and transposon proteins respectively.

**Figure 7:**
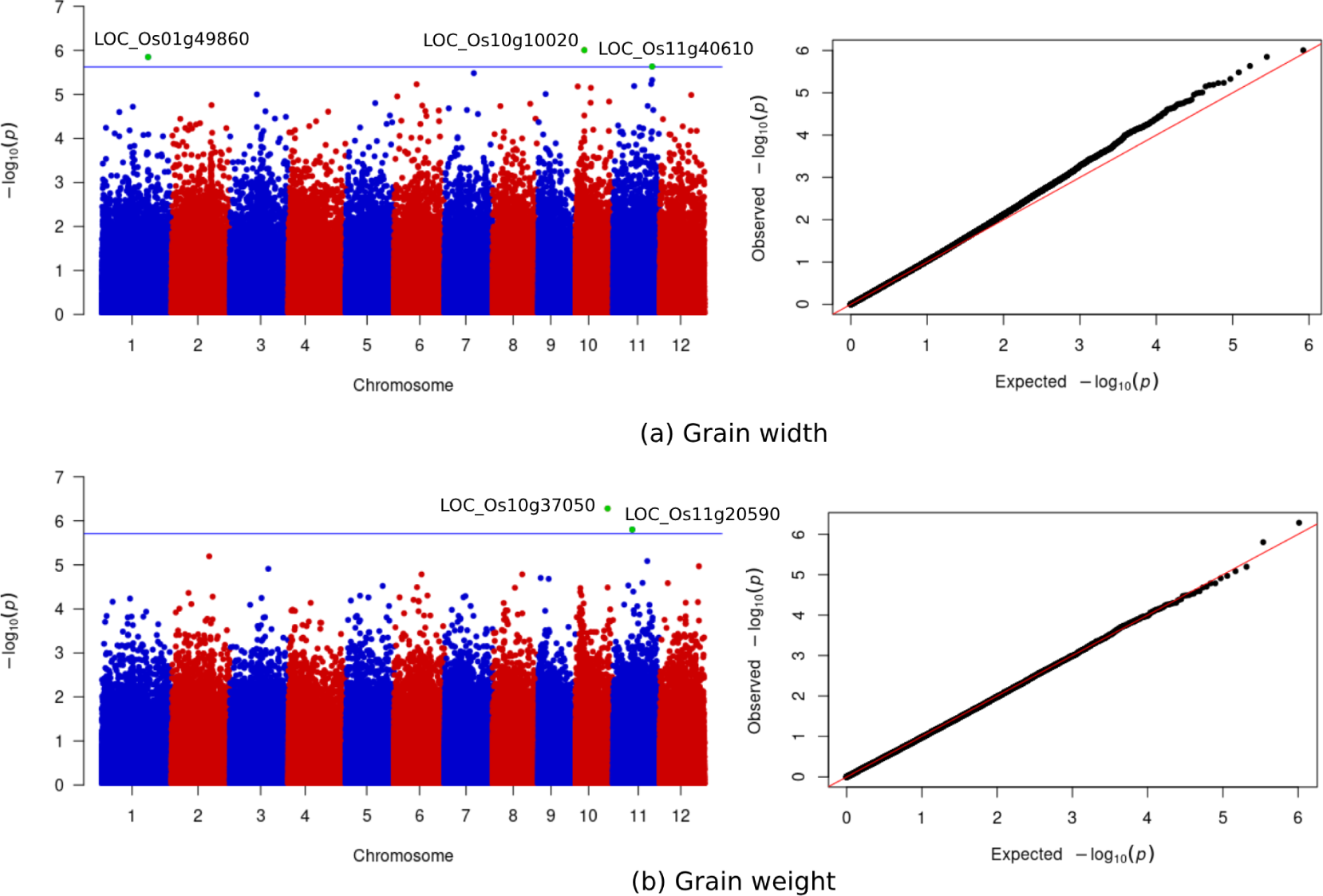
GWAS on grain width and weight using 168 rice varieties. We show the Manhattan plots and Q-Q plots for (a) grain width, and (b) grain weight.

The local Manhattan plots and LD heatmaps are presented in Fig. S5 in the supplementary material. LD analysis near the GWAS hit of chromosome 1 revealed zinc finger associated gene LOC Os01g49770. Zinc finger proteins are known to control cytokinin levels in the inflorescence meristem, which influences grain production [54]. We also observed genes nearby to the GWAS hit of chromosome 1 such as LOC Os01g49820, LOC Os01g49830, LOC Os01g49920, LOC Os01g49920, LOC Os01g49900 are highly expressed in inflorescence meristem. Another interesting gene we found in that region is LOC Os01g49870 which encodes a transposon.

Exploring the SNPs near the GWAS hit of chromosome 10, we found LOC Os10g09990 which encodes cytokinin-O-glucosyltransferase 3 (CGT) and previous GWA studies have shown that CGT genes have significance in grain maturity [55]. Other notable genes in that region are LOC Os10g10080 (exostosin family domain), LOC Os10g10040 (cytochrome P450), LOC Os10g10030 (OsWAK receptor-like protein kinase) as previous studies have shown their involvement in grain yield [56–58]. Moreover, we found a few retrotransposon proteins in this GWAS hit region (LOC Os10g09970, LOC Os10g09960, LOC Os10g09940, LOC Os10g09950, LOC Os10g09960, LOC Os10g09950).

In chromosome 11, near the GWAS hit site, we found LOC Os11g40550 (receptor kinase) and there is evidence that Mini Seed 2 (MIS2) gene encodes a receptor-like kinase which controls grain size and shape in rice [59]. Furthermore, LOC Os11g40680 encodes BTB (broad-complex, tramtrack, and bric-à-brac) protein complex that regulates the final grain size [60].

Ya-fang *et al.* (2015) [4] found chromosomes 1, 2, 3, 5, 6, 7, 10, 11 and 12 to harbor significant SNPs associated with GW. Yang *et al.* (2019) [9] identified various regions in chromosomes 4, 5, 6, 8, 9, and 11 to be associated with GW. Zhang *et al.* (2019) [7] discovered GS2 loci in chromosome 2 in close association with GW. Thus, our association study on GW shows similarities in chromosomal level (chromosomal region 1, 10, 11) with prior studies, in addition to revealing new loci-level associations.

#### 3.5.3 GWAS on grain weight (GWT)

Our study identified significantly associated SNPs harbored in chromosomes 10 and 11, suggesting two potential candidate genes LOC Os10g37050 and LOC Os11g20590 (Figure 7(b), and Table 2). These genes encode transposon proteins. Notably, we observed significant associations with transposon gene families for GL as well.

Our LD analysis (see Figure S6 in the supplementary material) near the GWAS hit of chromosome 11 revealed a cluster of genes such as LOC Os11g20510, LOC Os11g20610, LOC Os11g20520, LOC Os11g20570, and LOC Os11g20590, encoding retrotransposon and transposon proteins.

Our study identified gene LOC Os10g36924 near the GWAS hit on chromosome 10, which encodes an aquaporin protein. Aquaporins play crucial roles in water transport and plant physiological processes. While direct links to grain weight are not evident, aquaporins’ involvement in water transport suggests a potential indirect influence on grain weight by regulating plant hydration and nutrient uptake [61]. Additionally, LOC Os10g37034 encodes a cytochrome P450 protein, known to impact grain development [57]. We also noted the presence of a cytochrome gene (LOC Os10g10040) near the GWAS hit on chromosome 10 for the grain width trait.

As was observed for grain length and grain width, our association study on grain weight presents some congruent results with respect to the previous studies [62, 63], in addition to revealing new chromosomal regions associated with grain weight.

### 3.6 Results on leaf and panicle traits

We performed association studies on four other yield-related traits: panicle length and three leaf traits (leaf length (LL), leaf width (LW) and leaf angle (LA)). The Manhattan and Q-Q plots on these traits are shown in Fig. 8. The LD heatmap and local Manhattan plots are shown in Figures S7-S10 in the supplementary material.

**Figure 8:**
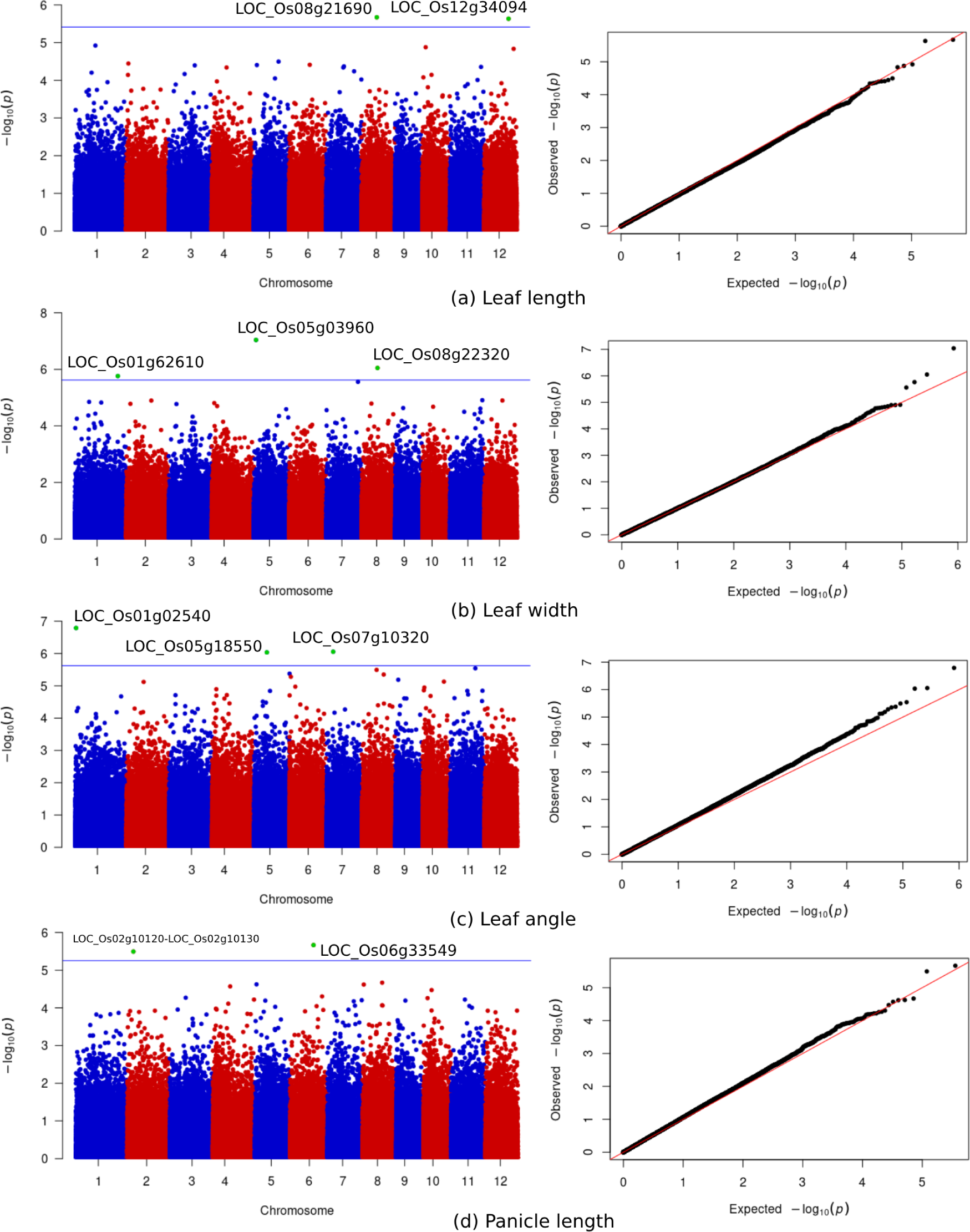
GWAS on leaf and panicle traits. We show the Manhattan and Q-Q plots for LL, LW, LA, and PL.

We find that two SNPs in chromosomes 8 and 12 are significantly associated with LL. Regarding LW, we see that three SNPs exceed the threshold line, located on chromosomes 1, 5, and 8 respectively. Three SNPs, harbored in chromosomes 1, 5, and 7, are observed to be significantly associated with LA. Finally, two SNPs in chromosomes 2 and 6 are found to be associated with PL. Details of the GWAS hits for these traits are shown in Table 2, SNPs that are in LD with the GWAS hits are in the supplementary data files (SM7-SM10).

## 4 Conclusions

Production of high-yield strains is crucial for meeting the continuously increasing food demand of the world population. Hybrid rice has been an effective way to meet this ever increasing food demand in Bangladesh. Despite a significant success in rice production, Bangladesh faces many challenges in the agricultural sector as it is becoming more densely populated day by day. In addition, climate change impacts like temperature rise, uncertain weather, prolonged dry season, irregular rainfall, frequent cyclones, sea-level rise, floods etc. are already being felt in Bangladesh. Therefore, understanding the genetic architecture underlying yield related traits as well as the impact of environmental factors are fundamental to the advancement of rice cultivars as the performance of various rice varieties vary with varying conditions of cultivation [64].

We presented genome-wide associated studies on 7 important yield-related traits using 183 rice varieties in Bangladesh. These GWASs are especially targeted for the Bangladeshi rice varieties, and thus consider the interactions between genetic variations underlying yield-related traits and the particular ecological environment of Bangladesh. The GWA studies for grain related traits (GL, GW, and GWT) yielded significant insights into the genetic determinants of these important agronomic traits. For grain length, significantly associted SNPs were identified on chromosomes 3, 7, and 10, corresponding to genes encoding retrotransposon proteins, known for their role in genome evolution. Similarly, GWAS for grain width revealed significant SNPs on chromosomes 1, 10, and 11, with candidate genes including an flowering and transposon proteins. The LD analysis near the GWAS hits provided further insights into the potential regulatory mechanisms, with zinc finger associated genes, highly expressed genes in inflorescence meristem, and BTB protein complex identified as candidates. Moreover, for grain weight, significant SNPs were found on chromosomes 10 and 11, with candidate genes encoding transposon proteins and aquaporin and cytochrome P450 proteins. Notably, the involvement of transposon gene families was observed across all three traits, suggesting a potential shared genetic basis.

This result could partially explain the genetic basis of correlation among the three grain-related traits (as demonstrated in Fig. 1), and provide useful information for genetic improvement of these traits by marker-assisted selection (MAS) [65–67]. As we discussed in our results section, there is discordance among the significant loci identified by various association studies performed on the same traits. Differences in sample sizes and various types of rice varieties considered in different studies may be attributed to this disagreement. Another crucial factor is the ecological environment where the considered rice varieties were grown. Therefore, this study advances the state-of-the-art in rice research in Bangladesh. However, this study is limited in scope, and can be extended in various directions. We have leveraged the data from the 3K RGP project. Future studies need to collect rice materials, planted in various regions of Bangladesh under adverse ecological conditions to better elucidate the impact of specific environmental factors in genotype-phenotype association. Follow-up studies also need to investigate the candidate genes through functional genomics approach [1, 68]. This study is limited to 7 yield-related traits. However, more information will be gained through GWAS of rice landraces as additional phenotypes are evaluated, especially the ones that are related to the adverse ecological environments of Bangladesh. To name a few, tolerance to prolonged flood, submergence, salinity, drought and cold are special features for various rice varieties in Bangladesh. As such, future studies need to sample a larger number of broadly representative varieties with special traits. For example, Rayada – a distinctive group of deepwater rice, totally endemic to certain area of Bangladesh and have multiple physiological features distinctly different from typical deepwater rice – could be potential resources of abiotic stress tolerance traits like flood, cold and drought [69–71]. Thus, we believe that this study will stimulate related future studies and will help identify beneficial genetic variations – which will enable the agricultural scientists to direct their efforts in developing elite varieties with desirable genetic compositions.

## Supplementary Materials

### Supplementary Material SM1: Additional figures and tables

**Figure S1:**
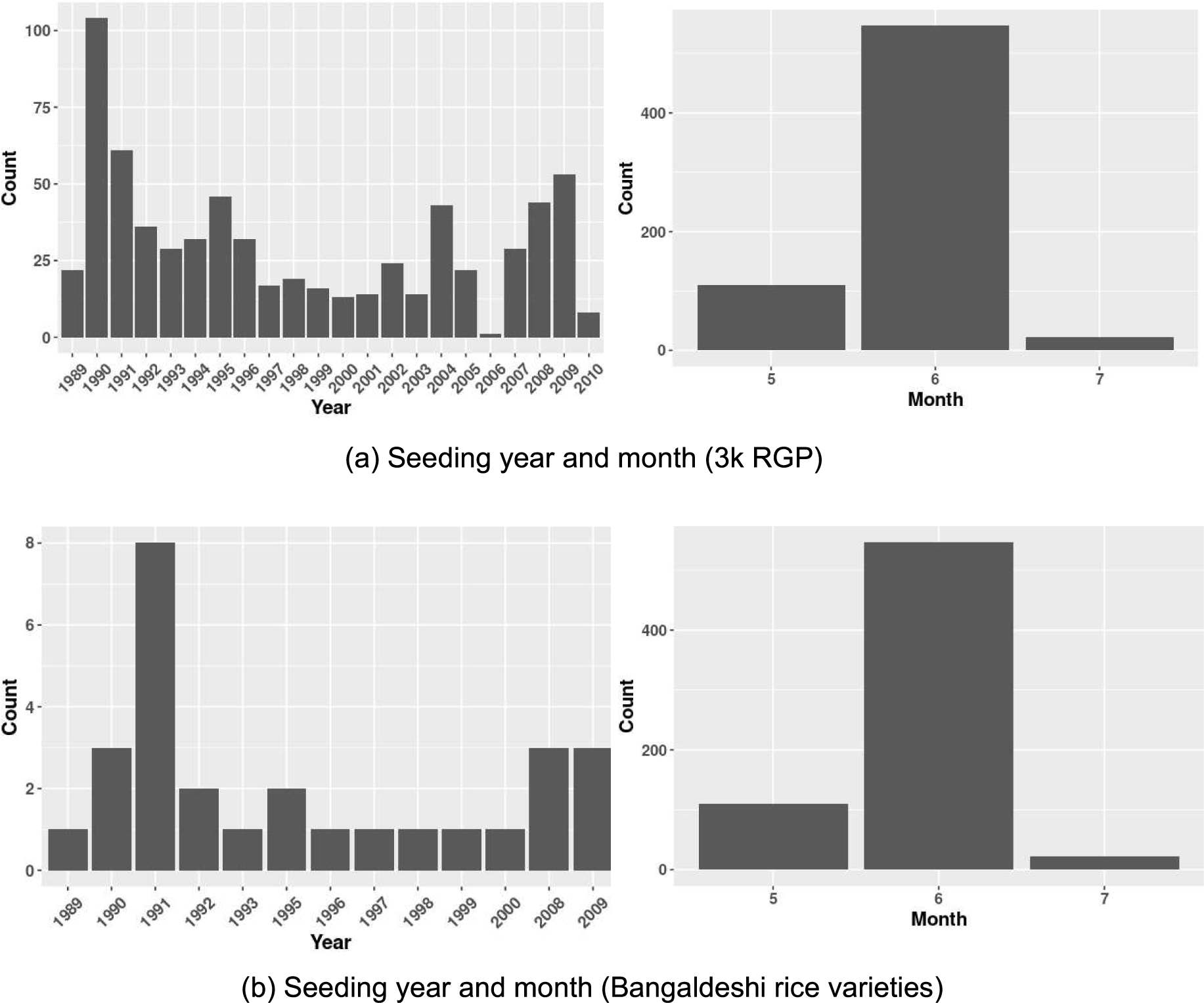
Seeding information of various rice varieties.

**Figure S2:**
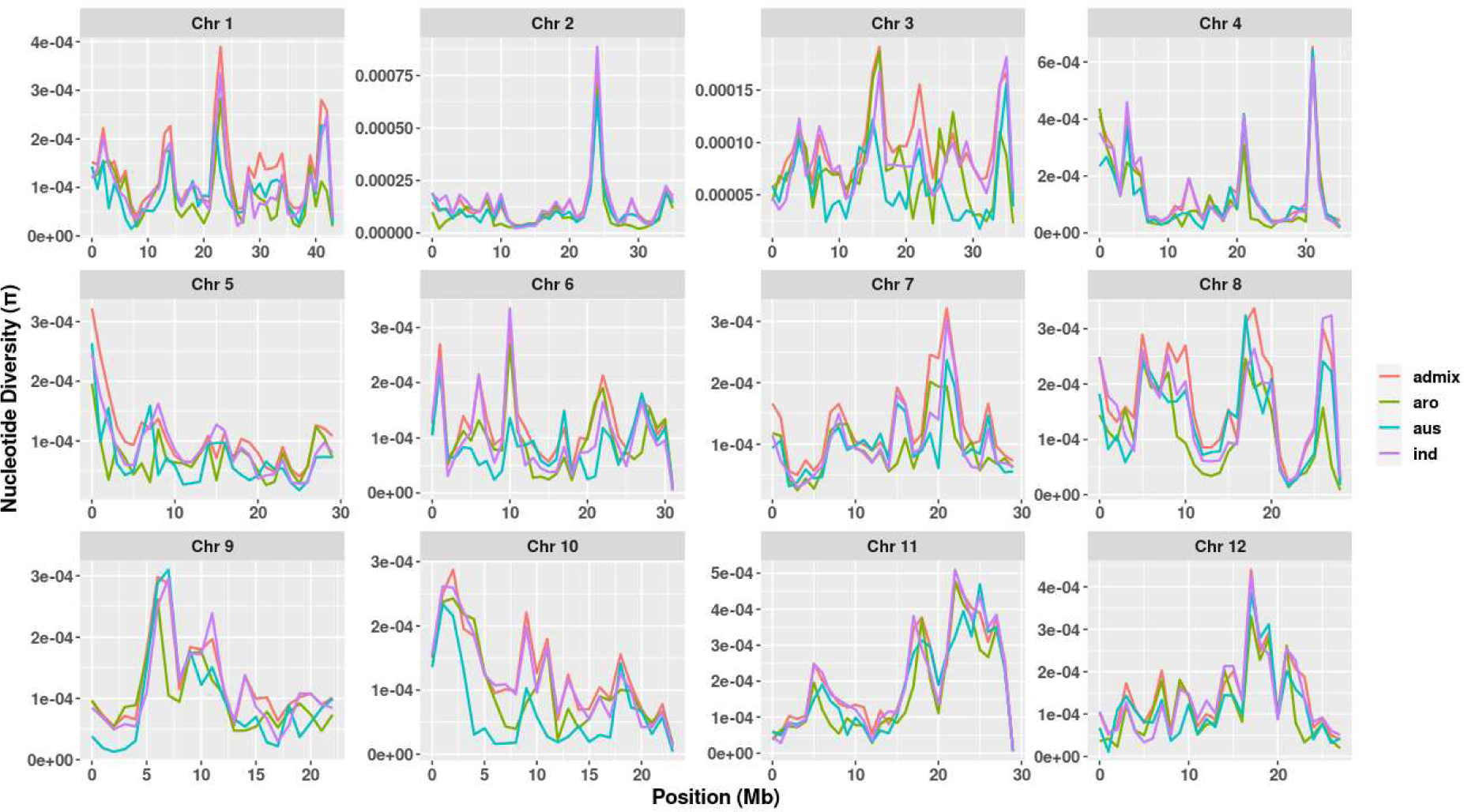
Nucleotide diversity (*π*) across chromosomes in Bangladeshi rice subpopulations.

**Figure S3:**
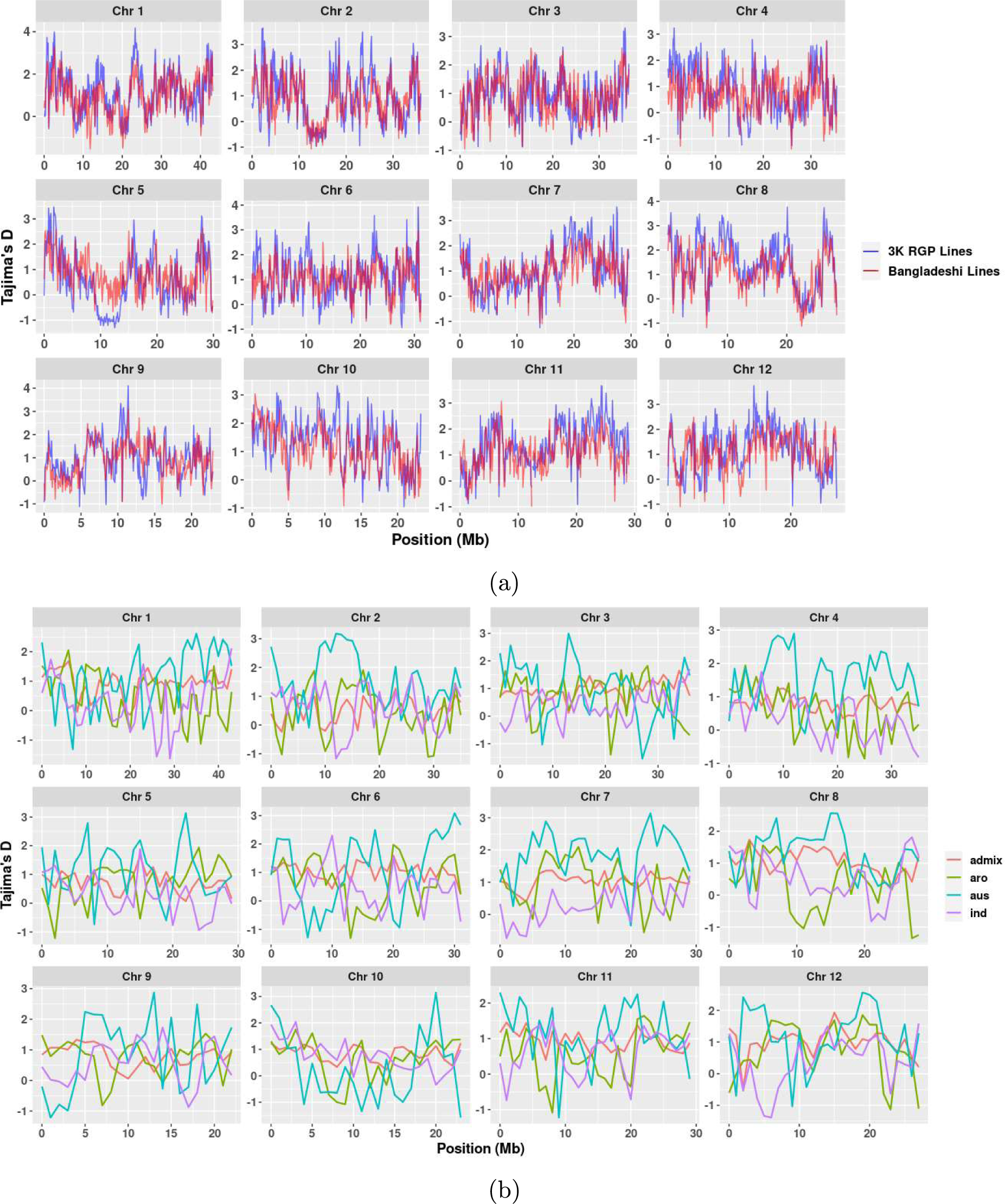
Tajima’s D distributions across chromosomes. (a) Comparing the distributions of 3K RGP and Bangladeshi rice lines, and (b) the distributions among the Bangladeshi lines.

**Figure S4:**
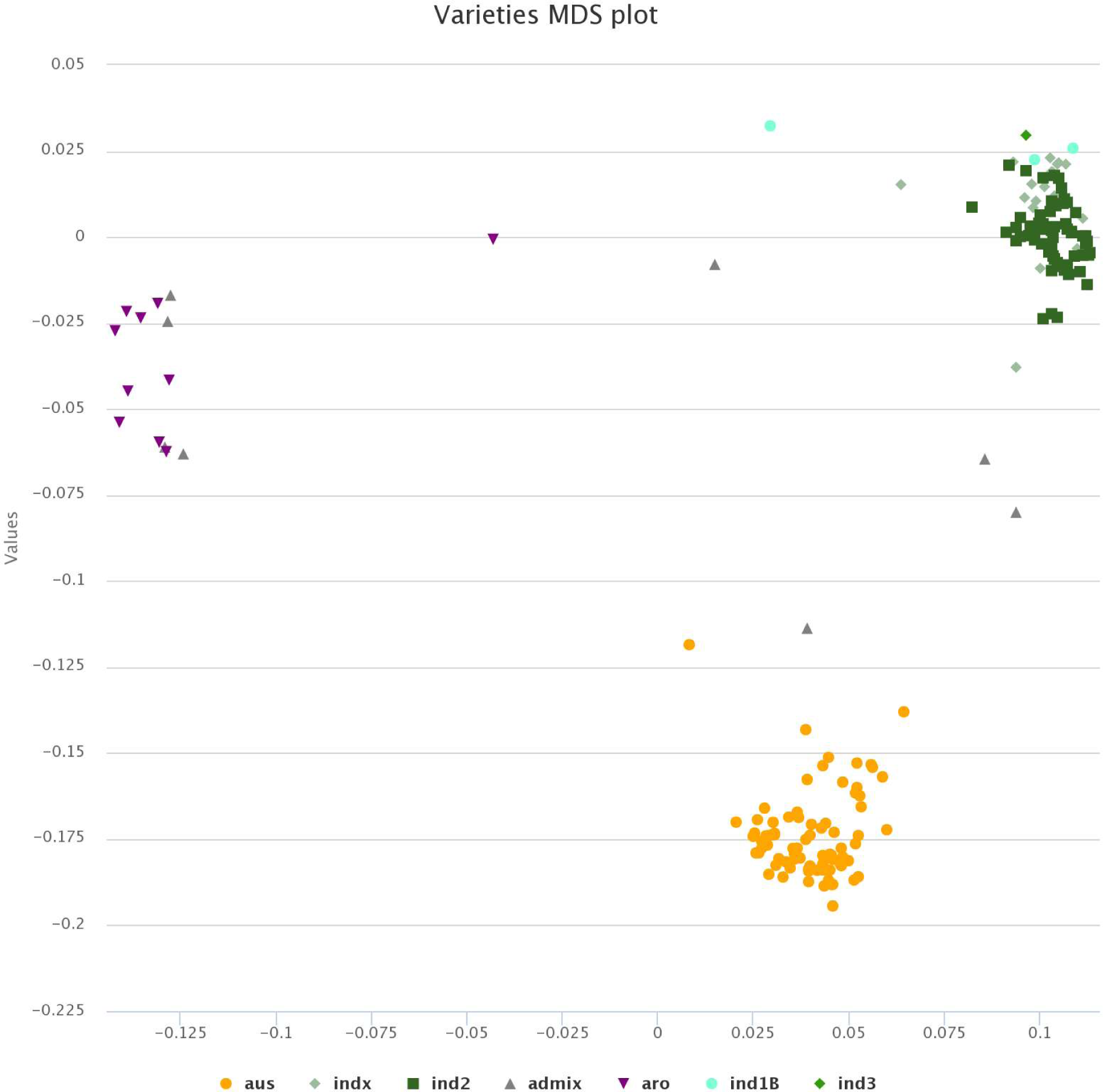
Genome stratification based on SNP markers using multidimensional scaling (MDS) plot for 183 rice varieties.

**Figure S5:**
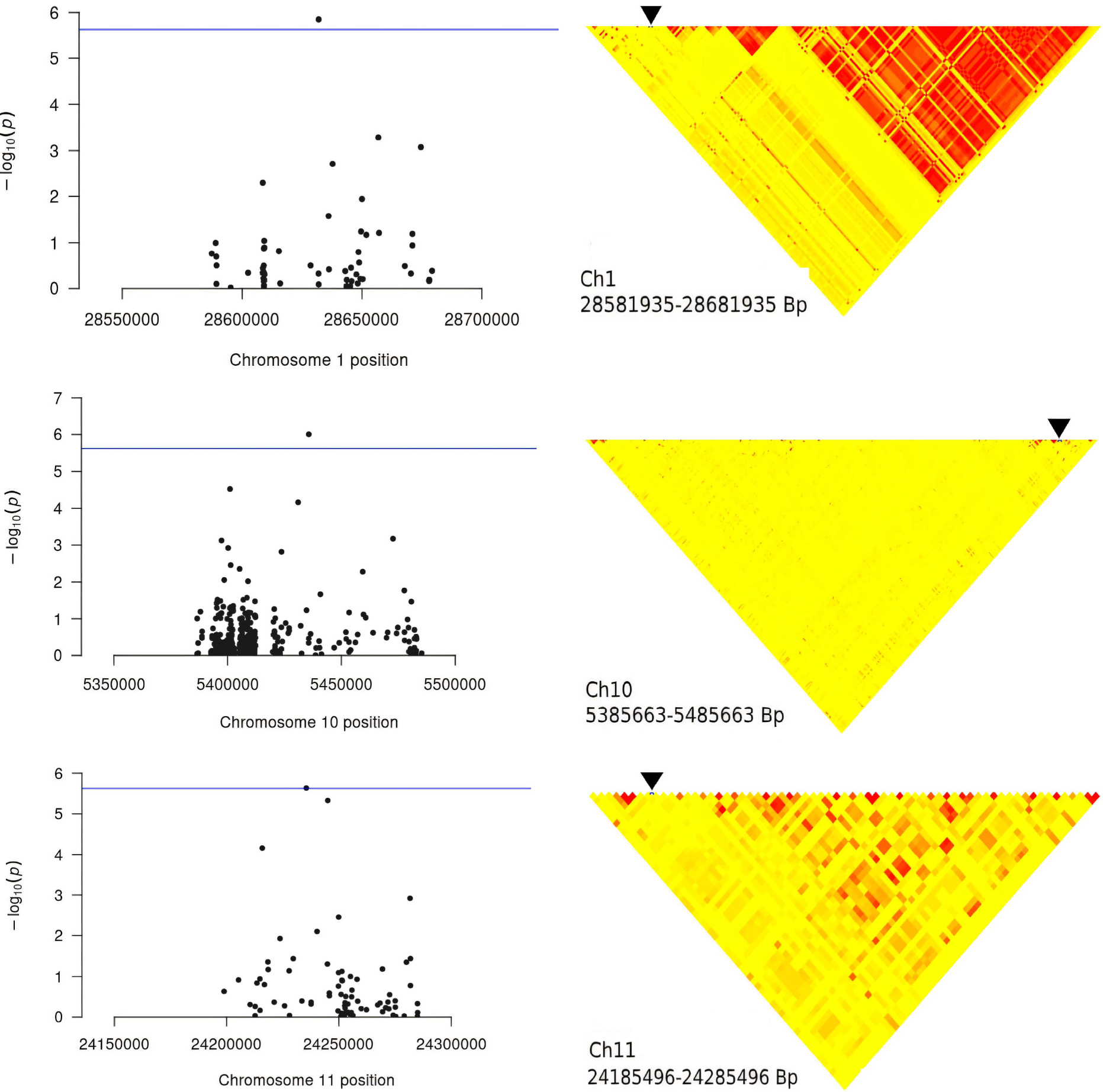
GWAS on grain width. We show the local Manhattan plots and LD heatmaps.

**Figure S6:**
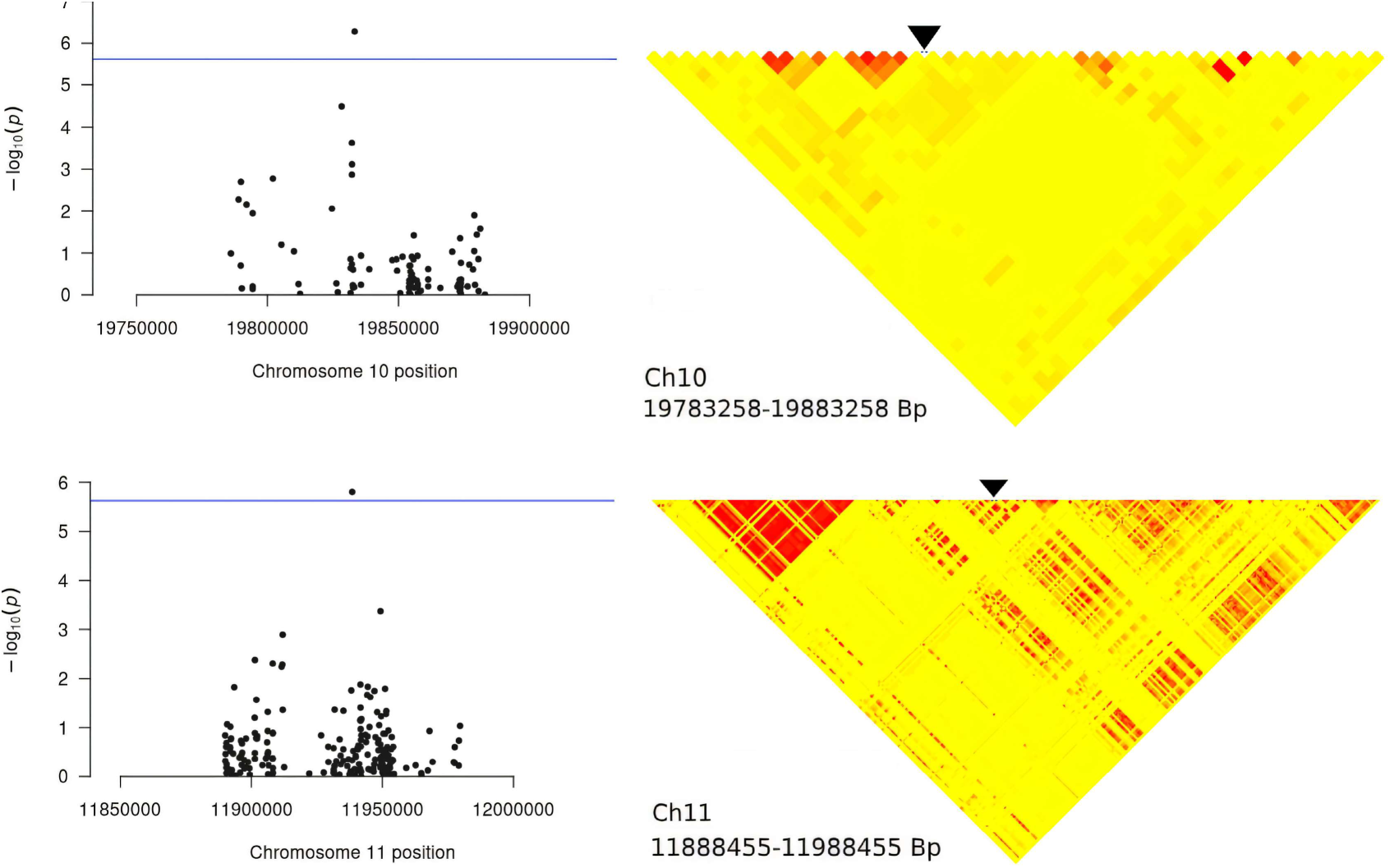
GWAS on grain weight. We show the local Manhattan plots and LD heatmaps.

**Figure S7:**
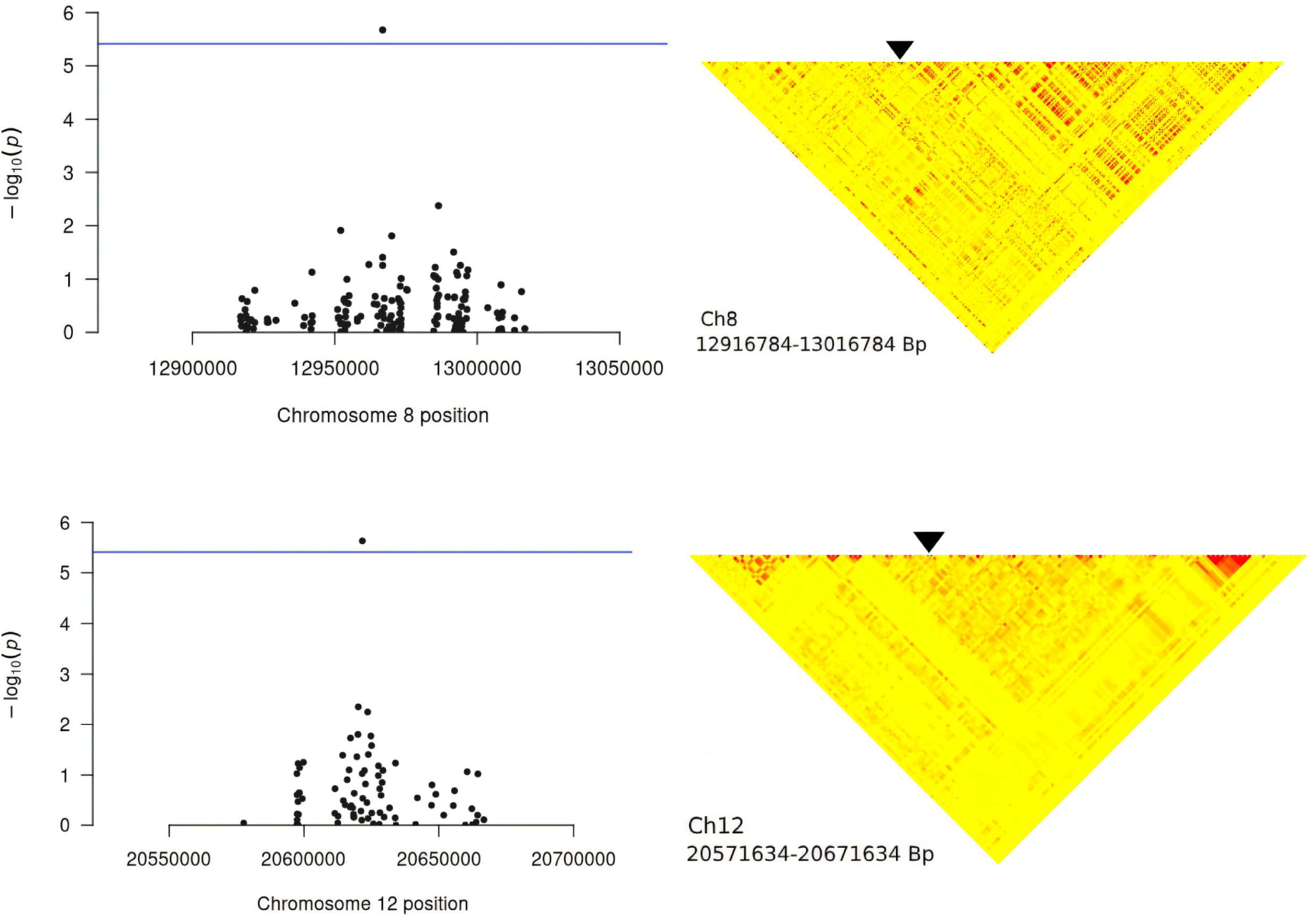
Local Manhattan plots and LD heatmaps for the GWAS on leaf length (LL).

**Figure S8:**
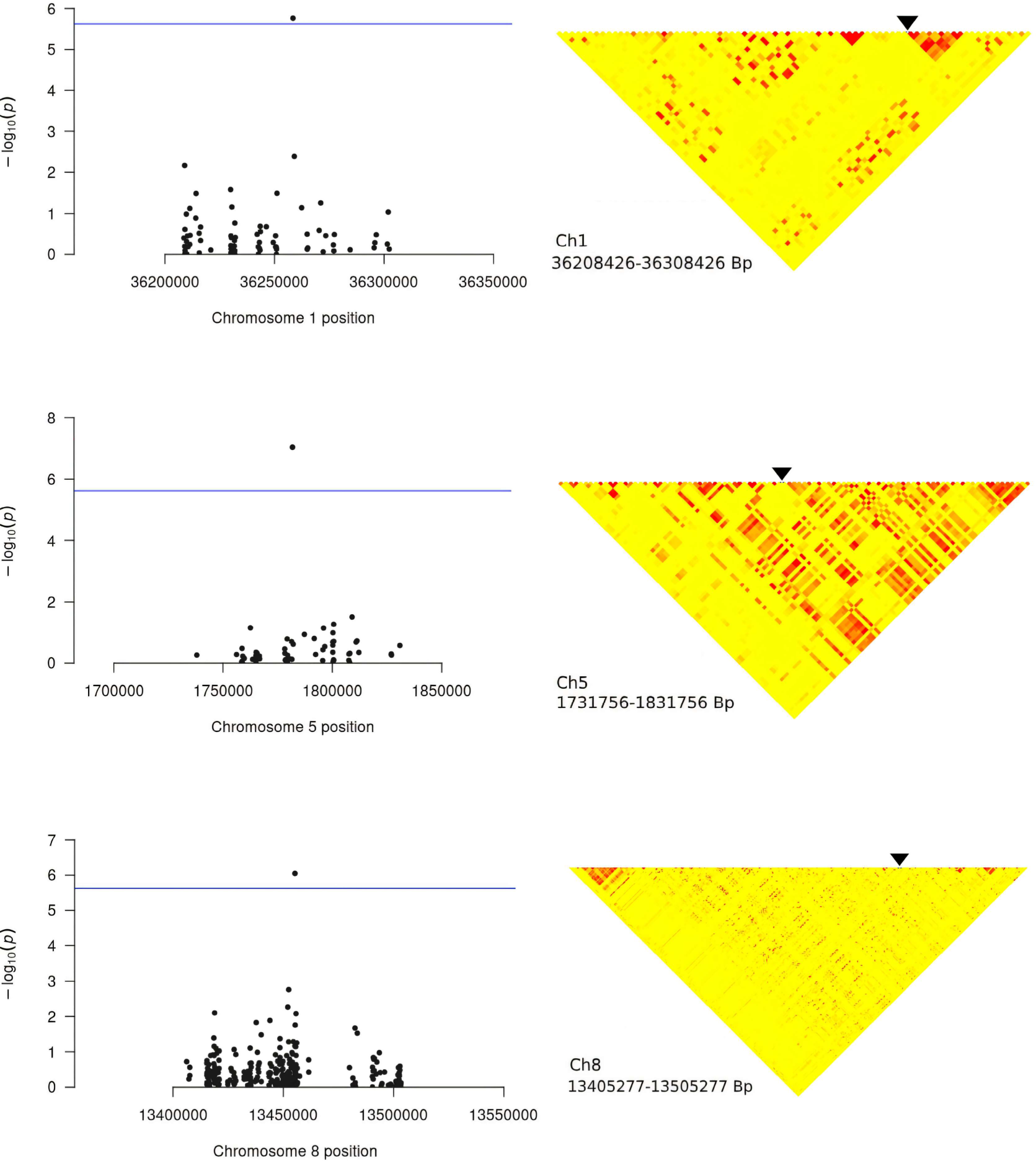
Local Manhattan plots and LD heatmaps for the GWAS on leaf width (LW).

**Figure S9:**
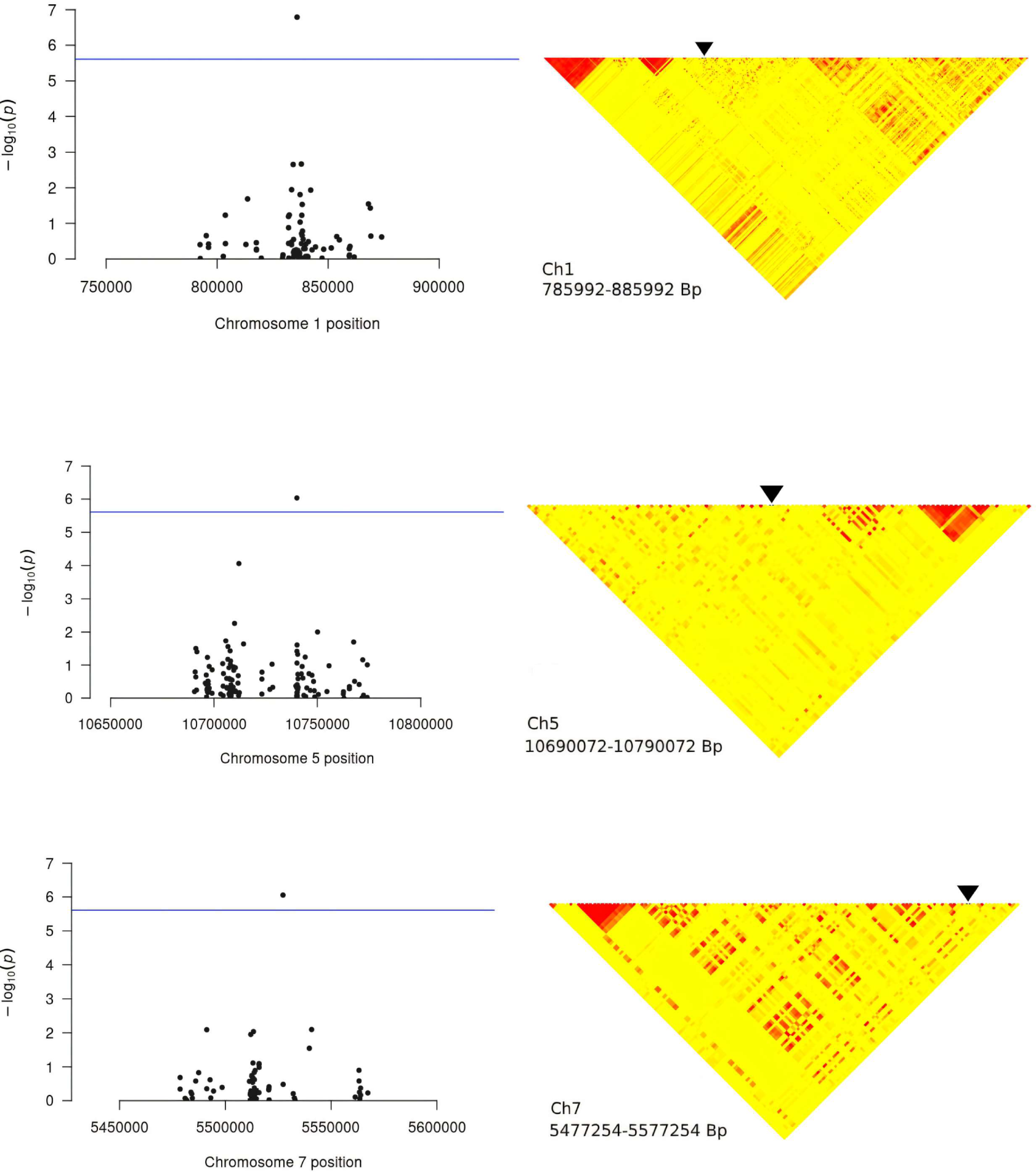
Local Manhattan plots and LD heatmaps for the GWAS on leaf angle (LA).

**Figure S10:**
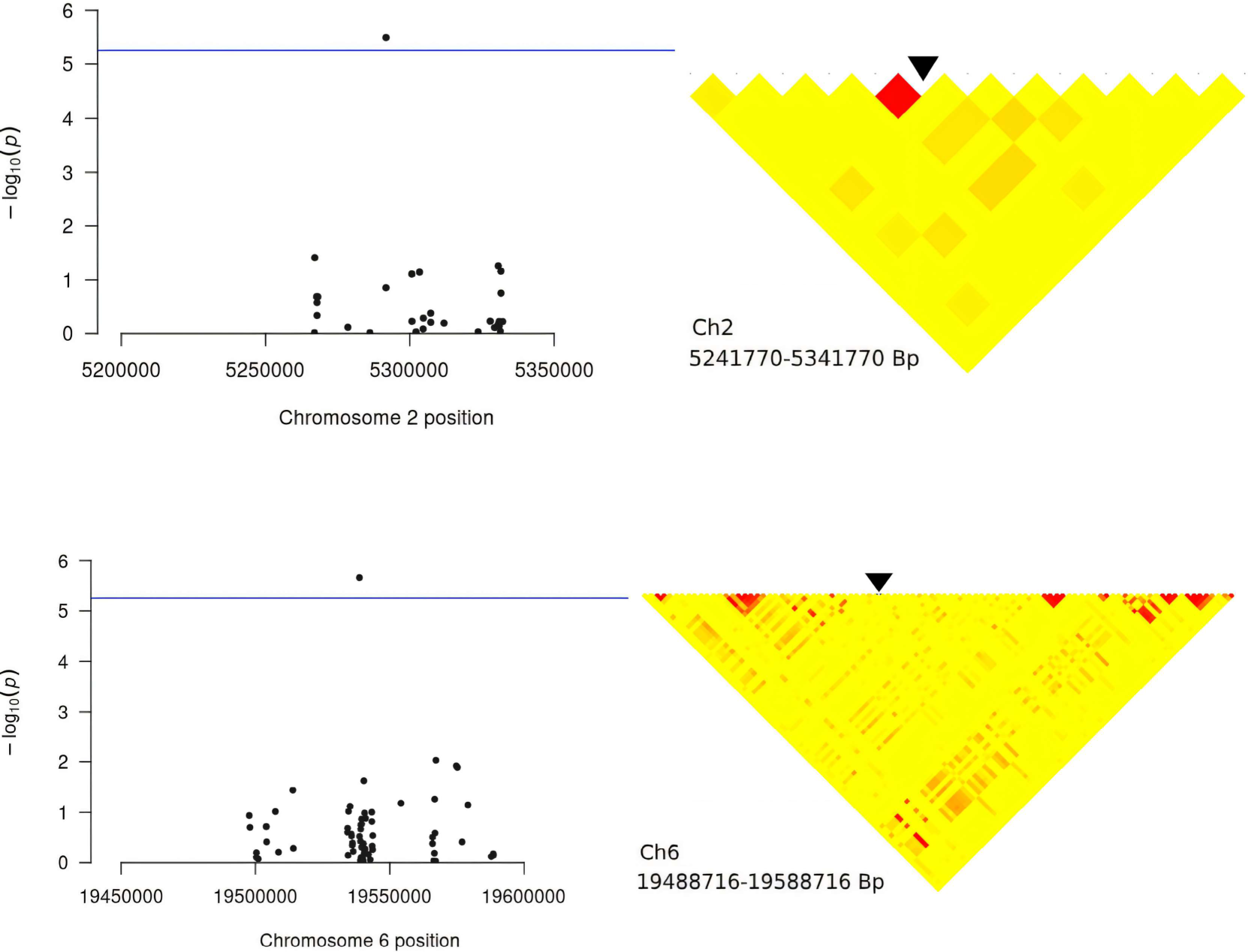
Local Manhattan plots and LD heatmaps for the GWAS on panicle length (PL).

**Figure S11:**
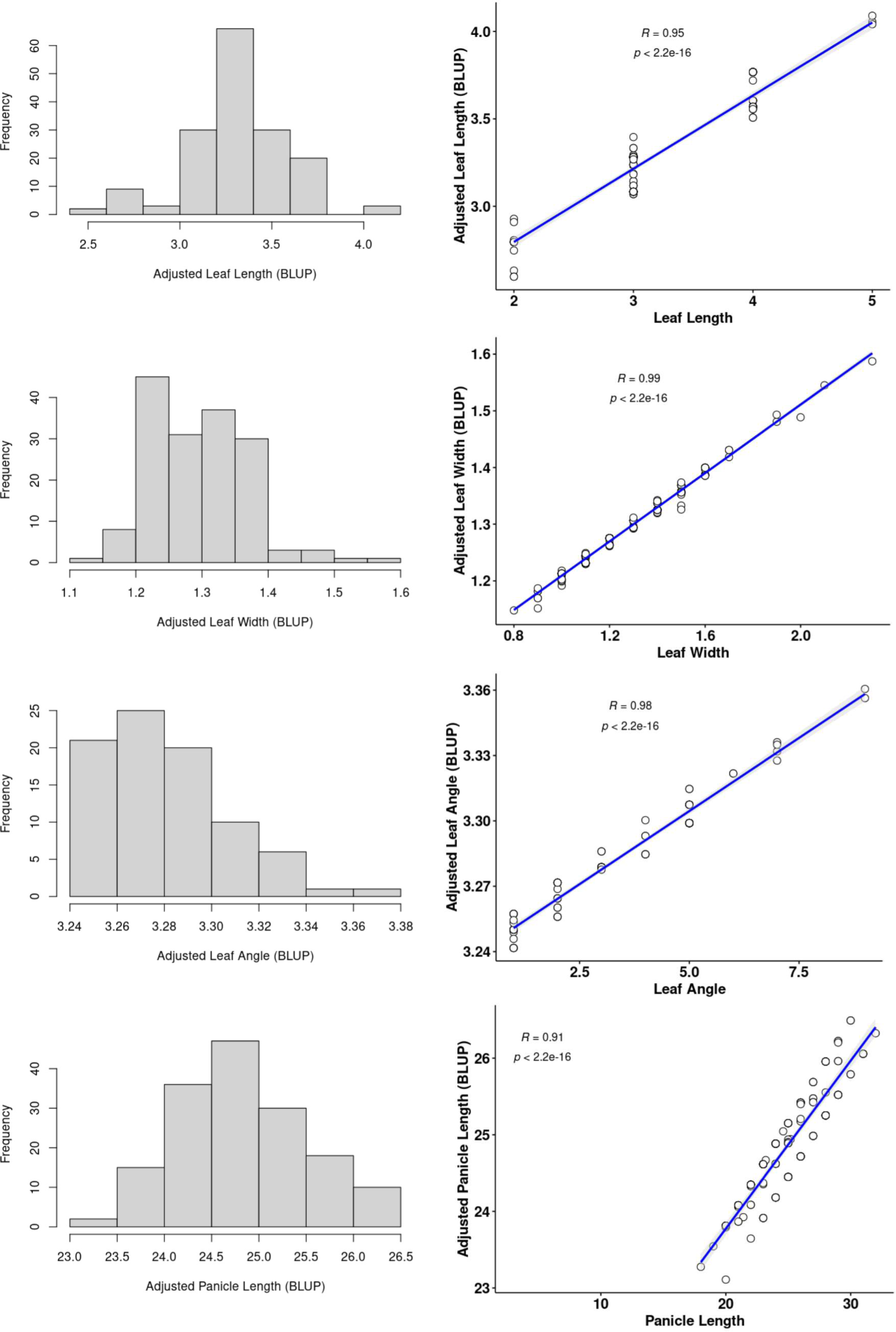
Distribution of BLUP-adjusted phenotypic values for leaf and panicle traits and the relationship of the raw values with the adjusted values.

**Figure S12:**
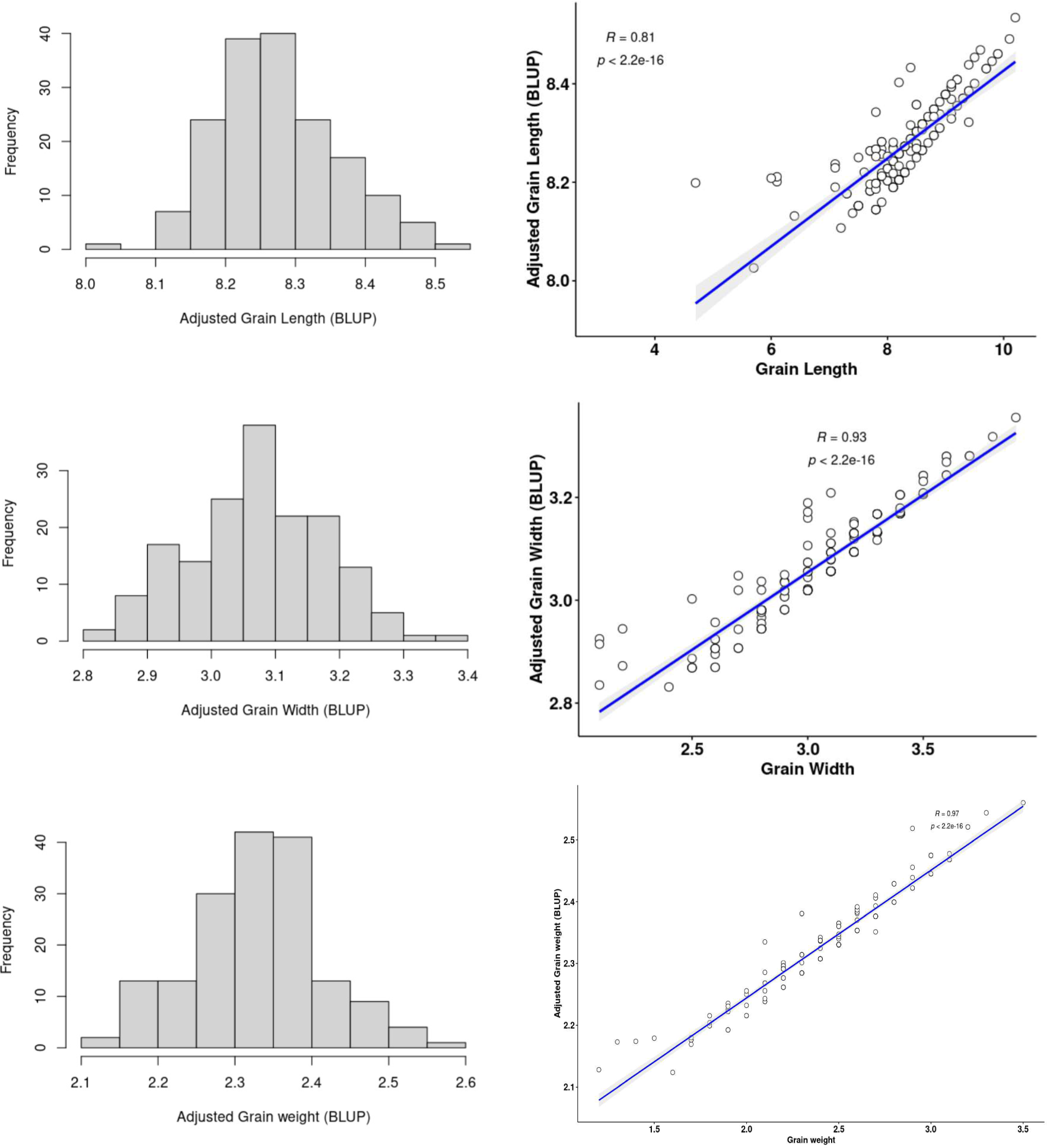
Distribution of BLUP-adjusted phenotypic values for grain shape related traits and the relationship of the raw values with the adjusted values.

**Data File SM2:** Detailed information on the 183 rice varieties considered in this study.

**Data File SM3:** Kinship matrix of the 183 rice varieties considered in this study.

**Data File SM4-SM10:** LD SNPs and their annotation for seven traits.

## Availability of data and material

The datasets analyzed in this paper are publicly available at snp-seek.irri.org.

## References

[1] Xuehui Huang, Tao Sang, Qiang Zhao, Qi Feng, Yan Zhao, Canyang Li, Chuanrang Zhu, Tingting Lu, Zhiwu Zhang, Meng Li, et al. Genome-wide association studies of 14 agronomic traits in rice landraces. Nature Genetics, 42(11):961, 2010.

[2] Xuehui Huang, Yan Zhao, Canyang Li, Ahong Wang, Qiang Zhao, Wenjun Li, Yunli Guo, Liuwei Deng, Chuanrang Zhu, Danlin Fan, et al. Genome-wide association study of flowering time and grain yield traits in a worldwide collection of rice germplasm. Nature Genetics, 44(1):32, 2012.

[3] Keyan Zhao, Chih-Wei Tung, Georgia C Eizenga, Mark H Wright, M Liakat Ali, Adam H Price, Gareth J Norton, M Rafiqul Islam, Andy Reynolds, Jason Mezey, et al. Genome-wide association mapping reveals a rich genetic architecture of complex traits in oryza sativa. Nature Communications, 2(1):1–10, 2011.

[4] Zhang Ya-fang, MA Yu-yin, Chen Zong-xiang, ZOU Jie, Chen Tian-xiao, LI Qianqian, Pan Xue-biao, and Zuo Shi-min. Genome-wide association studies reveal new genetic targets for five panicle traits of international rice varieties. Rice Science, 22(5):217–226, 2015.

[5] Wanneng Yang, Zilong Guo, Chenglong Huang, Lingfeng Duan, Guoxing Chen, Ni Jiang, Wei Fang, Hui Feng, Weibo Xie, Xingming Lian, et al. Combining highthroughput phenotyping and genome-wide association studies to reveal natural genetic variation in rice. Nature Communications, 5:5087, 2014.

[6] Filippo Biscarini, Paolo Cozzi, Laura Casella, Paolo Riccardi, Alessandra Vattari, Gabriele Orasen, Rosaria Perrini, Gianni Tacconi, Alessandro Tondelli, Chiara Biselli, et al. Genome-wide association study for traits related to plant and grain morphology, and root architecture in temperate rice accessions. PloS One, 11(5), 2016.

[7] Peng Zhang, Kaizhen Zhong, Zhengzheng Zhong, and Hanhua Tong. Genome-wide association study of important agronomic traits within a core collection of rice (Oryza sativa l.). BMC Plant Biology, 19(1):259, 2019.

[8] Xiao-ling LI, Yong-gen Lu, Jin-quan Li, Hai-ming Xu, and Muhammad Qasim Shahihd. Strategies on sample size determination and qualitative and quantitative traits integration to construct core collection of rice (oryza sativa). Rice Science, 18(1):46–55, 2011.

[9] LV Yang, Wang Yueying, Noushin Jahan, Hu Haitao, Chen Ping, Shang Lianguang, Lin Haiyan, Dong Guojun, Hu Jiang, Gao Zhenyu, et al. Genome-wide association analysis and allelic mining of grain shape-related traits in rice. Rice Science, 26(6):384–392, 2019.

[10] Xiaosong Ma, Fangjun Feng, Yu Zhang, Ibrahim Eid Elesawi, Kai Xu, Tianfei Li, Hanwei Mei, Hongyan Liu, Ningning Gao, Chunli Chen, et al. A novel rice grain size gene ossnb was identified by genome-wide association study in natural population. PLoS Genetics, 15(5):e1008191, 2019.

[11] Wanneng Yang, Zilong Guo, Chenglong Huang, Ke Wang, Ni Jiang, Hui Feng, Guoxing Chen, Qian Liu, and Lizhong Xiong. Genome-wide association study of rice (oryza sativa l.) leaf traits with a high-throughput leaf scorer. Journal of Experimental Botany, 66(18):5605–5615, 2015.

[12] 3000 Rice Genomes Project. The 3,000 rice genomes project. GigaScience, 3(1):2047–217X, 2014.

[13] Nickolai Alexandrov, Shuaishuai Tai, Wensheng Wang, Locedie Mansueto, Kevin Palis, Roven Rommel Fuentes, Victor Jun Ulat, Dmytro Chebotarov, Gengyun Zhang, Zhikang Li, et al. Snp-seek database of snps derived from 3000 rice genomes. Nucleic Acids Research, 43(D1):D1023–D1027, 2015.

[14] Locedie Mansueto, Roven Rommel Fuentes, Frances Nikki Borja, Jeffery Detras, Juan Miguel Abriol-Santos, Dmytro Chebotarov, Millicent Sanciangco, Kevin Palis, Dario Copetti, Alexandre Poliakov, et al. Rice snp-seek database update: new snps, indels, and queries. Nucleic Acids Research, 45(D1):D1075–D1081, 2017.

[15] Rongyu Huang, Liangrong Jiang, Jingsheng Zheng, Tiansheng Wang, Houcong Wang, Yumin Huang, and Zonglie Hong. Genetic bases of rice grain shape: so many genes, so little known. Trends in Plant Science, 18(4):218–226, 2013.

[16] Mayuko Ikeda, Kotaro Miura, Koichiro Aya, Hidemi Kitano, and Makoto Matsuoka. Genes offering the potential for designing yield-related traits in rice. Current Opinion in Plant Biology, 16(2):213–220, 2013.

[17] HX Huang and Q Qian. Progress in genetic research of rice grain shape and breeding achievements of long-grain shape and good quality japonica rice. Chin J Rice Sci, 31(6):665–672, 2017.

[18] Giang Thi Hoang, Pascal Gantet, Kien Huu Nguyen, Nhung Thi Phuong Phung, Loan Thi Ha, Tuan Thanh Nguyen, Michel Lebrun, Brigitte Courtois, and Xuan Hoi Pham. Genome-wide association mapping of leaf mass traits in a vietnamese rice landrace panel. PloS One, 14(7), 2019.

[19] Peng Wang, Guilin Zhou, Huihui Yu, and Sibin Yu. Fine mapping a major qtl for flag leaf size and yield-related traits in rice. Theoretical and Applied Genetics, 123(8):1319–1330, 2011.

[20] Hirokazu Tsukaya. Leaf shape: genetic controls and environmental factors. International Journal of Developmental Biology, 49(5-6):547–555, 2004.

[21] José Manuel Pérez-Pérez, David Esteve-Bruna, and José Luis Micol. Qtl analysis of leaf architecture. Journal of Plant Research, 123(1):15–23, 2010.

[22] Baoyan Jia, Xinhua Zhao, Yang Qin, Muhammad Irfan, Tae-heon Kim, Bolun Wang, Shu Wang, and Jae Keun Sohn. Quantitative trait loci mapping of panicle traits in rice. Molecular Biology Research Communications, 8(1):9, 2019.

[23] International Rice Genebank Operations Manual. https://cropgenebank.sgrp.cgiar.org/images/file/learning_space/IRRI_genebank_manual.pdf. Accessed: 2021-07-01.

[24] Wensheng Wang, Ramil Mauleon, Zhiqiang Hu, Dmytro Chebotarov, Shuaishuai Tai, Zhichao Wu, Min Li, Tianqing Zheng, Roven Rommel Fuentes, Fan Zhang, et al. Genomic variation in 3,010 diverse accessions of asian cultivated rice. Nature, 557(7703):43–49, 2018.

[25] 000 Rice Genomes Project 3. The 3,000 rice genomes project. Gigascience, 3(1):2047–217X, 2014.

[26] R R Core Team, et al. R: A language and environment for statistical computing. 2013.

[27] Philip Dixon. Vegan, a package of r functions for community ecology. Journal of vegetation science, 14(6):927–930, 2003.

[28] Douglas Bates, Martin Mächler, Ben Bolker, and Steve Walker. Fitting linear mixedeffects models using lme4. arXiv preprint arXiv:1406.5823, 2014.

[29] Shaun Purcell, Benjamin Neale, Kathe Todd-Brown, Lori Thomas, Manuel AR Ferreira, David Bender, Julian Maller, Pamela Sklar, Paul IW De Bakker, Mark J Daly, et al. Plink: a tool set for whole-genome association and population-based linkage analyses. The American Journal of Human Genetics, 81(3):559–575, 2007.

[30] Petr Danecek, Adam Auton, Goncalo Abecasis, Cornelis A Albers, Eric Banks, Mark A DePristo, Robert E Handsaker, Gerton Lunter, Gabor T Marth, Stephen T Sherry, et al. The variant call format and vcftools. Bioinformatics, 27(15):2156–2158, 2011.

[31] Camilla J Williams, Zhixiu Li, Nicholas Harvey, Rodney A Lea, Brendon J Gurd, Jacob T Bonafiglia, Ioannis Papadimitriou, Macsue Jacques, Ilaria Croci, Dorthe Stensvold, et al. Genome wide association study of response to interval and continuous exercise training: the predict-hiit study. Journal of biomedical science, 28(1):37, 2021.

[32] Qing Lu, Mengchen Zhang, Xiaojun Niu, Shan Wang, Qun Xu, Yue Feng, Caihong Wang, Hongzhong Deng, Xiaoping Yuan, Hanyong Yu, et al. Genetic variation and association mapping for 12 agronomic traits in indica rice. BMC genomics, 16:1–17, 2015.

[33] Jeffrey C Barrett, B Fry, JDMJ Maller, and Mark J Daly. Haploview: analysis and visualization of ld and haplotype maps. Bioinformatics, 21(2):263–265, 2005.

[34] Ji-Hyung Shin, Sigal Blay, Brad McNeney, and Jinko Graham. Ldheatmap: an r function for graphical display of pairwise linkage disequilibria between single nucleotide polymorphisms. Journal of statistical software, 16:1–9, 2006.

[35] Yoshihiro Kawahara, Melissa de la Bastide, John P Hamilton, Hiroyuki Kanamori, W Richard McCombie, Shu Ouyang, David C Schwartz, Tsuyoshi Tanaka, Jianzhong Wu, Shiguo Zhou, et al. Improvement of the oryza sativa nipponbare reference genome using next generation sequence and optical map data. Rice, 6(1):4, 2013.

[36] Hadley Wickham, Mara Averick, Jennifer Bryan, Winston Chang, Lucy D’Agostino McGowan, Romain François, Garrett Grolemund, Alex Hayes, Lionel Henry, Jim Hester, et al. Welcome to the tidyverse. Journal of open source software, 4(43):1686, 2019.

[37] Subhas Chandra Roy and Pankaj Shil. Assessment of genetic heritability in rice breeding lines based on morphological traits and caryopsis ultrastructure. Scientific reports, 10(1):7830, 2020.

[38] S Widyayanti, H Purwaningsih, et al. Genetic diversity of local red rice cultivars collections of yogyakarta aiat, indonesia based on morphological character. In IOP Conference Series: Earth and Environmental Science, volume 482, page 012040. IOP Publishing, 2020.

[39] Jeffrey B Endelman and Jean-Luc Jannink. Shrinkage estimation of the realized relationship matrix. G3: Genes— Genomes— Genetics, 2(11):1405–1413, 2012.

[40] Peter J Bradbury, Zhiwu Zhang, Dallas E Kroon, Terry M Casstevens, Yogesh Ramdoss, and Edward S Buckler. Tassel: software for association mapping of complex traits in diverse samples. Bioinformatics, 23(19):2633–2635, 2007.

[41] Eleonora Porcu, Sina Rüeger, Kaido Lepik, Federico A Santoni, Alexandre Reymond, and Zoltán Kutalik. Mendelian randomization integrating gwas and eqtl data reveals genetic determinants of complex and clinical traits. Nature communications, 10(1):3300, 2019.

[42] Simon Orozco-Arias, Gustavo Isaza, and Romain Guyot. Retrotransposons in plant genomes: structure, identification, and classification through bioinformatics and machine learning. International journal of molecular sciences, 20(15):3837, 2019.

[43] Pradeep K Papolu, Muthusamy Ramakrishnan, Sileesh Mullasseri, Ruslan Kalendar, Qiang Wei, Long Zou, Zishan Ahmad, Kunnummal Kurungara Vinod, Ping Yang, Mingbing Zhou, et al. Retrotransposons: How the continuous evolutionary front shapes plant genomes for response to heat stress. Frontiers in plant science, 13:1064847, 2022.

[44] Roland Akakpo, Marie-Christine Carpentier, Yue Ie Hsing, and Olivier Panaud. The impact of transposable elements on the structure, evolution and function of the rice genome. New Phytologist, 226(1):44–49, 2020.

[45] Dong-Min Kim, Hyun-Sook Lee, Soo-Jin Kwon, Mark Edward Fabreag, Ju-Won Kang, Yeo-Tae Yun, Chong-Tae Chung, and Sang-Nag Ahn. High-density mapping of quantitative trait loci for grain-weight and spikelet number in rice. Rice, 7(1):1–11, 2014.

[46] Jing Jin, Lei Hua, Zuofeng Zhu, Lubin Tan, Xinhui Zhao, Weifeng Zhang, Fengxia Liu, Yongcai Fu, Hongwei Cai, Xianyou Sun, et al. Gad1 encodes a secreted peptide that regulates grain number, grain length, and awn development in rice domestication. The Plant Cell, 28(10):2453–2463, 2016.

[47] Luling Xiong, Yingyong Huang, Zupei Liu, Chen Li, Hang Yu, Muhammad Qasim Shahid, Yanhui Lin, Xiaoyi Qiao, Junyi Xiao, Julie E Gray, et al. Small epidermal patterning factor-like2 peptides regulate awn development in rice. Plant Physiology, 190(1):516–531, 2022.

[48] Kenji Kitamura, Mai Nakase, Hideki Tohda, and Kaoru Takegawa. The ubiquitin ligase ubr11 is essential for oligopeptide utilization in the fission yeast schizosaccharomyces pombe. Eukaryotic cell, 11(3):302–310, 2012.

[49] Xinghai Yang, Xiuzhong Xia, Yu Zeng, Baoxuan Nong, Zongqiong Zhang, Yanyan Wu, Qinglan Tian, Weiying Zeng, Ju Gao, Weiyong Zhou, et al. Genome-wide identification of the peptide transporter family in rice and analysis of the ptr expression modulation in two near-isogenic lines with different nitrogen use efficiency. BMC plant biology, 20:1–15, 2020.

[50] Noriko Takano-Kai, Hui Jiang, Takahiko Kubo, Megan Sweeney, Takashi Matsumoto, Hiroyuki Kanamori, Badri Padhukasahasram, Carlos Bustamante, Atsushi Yoshimura, Kazuyuki Doi, et al. Evolutionary history of gs3, a gene conferring grain length in rice. Genetics, 182(4):1323–1334, 2009.

[51] Rakesh Singh, Ashok Kumar Singh, Tilak Raj Sharma, Aqbal Singh, and Nagendra K Singh. Fine mapping of grain length qtls on chromosomes 1 and 7 in basmati rice (oryza sativa l.). Journal of Plant Biochemistry and Biotechnology, 21(2):157–166, 2012.

[52] Kashif Aslam and Muhammad Arif. Ssr analysis of chromosomes 3 and 7 of rice (oryza staiva l.) associated with grain length. Pak. J. Bot, 46(4):1363–1372, 2014.

[53] Tsutomu Ishimaru, Khin Thandar Hlaing, Ye Min Oo, Tin Mg Lwin, Kazuhiro Sasaki, Patrick D Lumanglas, Eliza-Vie M Simon, Tin Tin Myint, Aris Hairmansis, Untung Susanto, et al. An early-morning flowering trait in rice can enhance grain yield under heat stress field conditions at flowering stage. Field Crops Research, 277:108400, 2022.

[54] Shuyu Li, Bingran Zhao, Dingyang Yuan, Meijuan Duan, Qian Qian, Li Tang, Bao Wang, Xiaoqiang Liu, Jie Zhang, Jun Wang, et al. Rice zinc finger protein dst enhances grain production through controlling gn1a/osckx2 expression. Proceedings of the National Academy of Sciences, 110(8):3167–3172, 2013.

[55] Wadzani Palnam Dauda, Veerubommu Shanmugam, Aditya Tyagi, Amolkumar U Solanke, Vishesh Kumar, Subbaiyan Gopala Krishnan, Bishnu Maya Bashyal, and Rashmi Aggarwal. Genome-wide identification and characterisation of cytokinino-glucosyltransferase (cgt) genes of rice specific to potential pathogens. Plants, 11(7):917, 2022.

[56] Qilin Mu, Wenying Zhang, Yunbo Zhang, Haoliang Yan, Ke Liu, Tsutomu Matsui, Xiaohai Tian, and Pingfang Yang. itraq-based quantitative proteomics analysis on rice anther responding to high temperature. International journal of molecular sciences, 18(9):1811, 2017.

[57] Babar Usman, Gul Nawaz, Neng Zhao, Yaoguang Liu, and Rongbai Li. Generation of high yielding and fragrant rice (oryza sativa l.) lines by crispr/cas9 targeted mutagenesis of three homoeologs of cytochrome p450 gene family and osbadh2 and transcriptome and proteome profiling of revealed changes triggered by mutations. Plants, 9(6):788, 2020.

[58] Hao Chen, Yanyan Tang, Jianfeng Liu, Lubin Tan, Jiahuan Jiang, Mumu Wang, Zuofeng Zhu, Xianyou Sun, and Chuanqing Sun. Emergence of a novel chimeric gene underlying grain number in rice. Genetics, 205(2):993–1002, 2017.

[59] Yan Chun, Jingjing Fang, Syed Adeel Zafar, Jiangyuan Shang, Jinfeng Zhao, Shoujiang Yuan, and Xueyong Li. Mini seed 2 (mis2) encodes a receptor-like kinase that controls grain size and shape in rice. Rice, 13:1–17, 2020.

[60] Abdullah Shalmani, Uzair Ullah, Li Tai, Ran Zhang, Xiu-Qing Jing, Izhar Muhammad, Nadeem Bhanbhro, Wen-Ting Liu, Wen-Qiang Li, and Kun-Ming Chen. Osbbx19-osbtb97/osbbx11 module regulates spikelet development and yield production in rice. Plant Science, 334:111779, 2023.

[61] Hidehiro Hayashi, Junko Ishikawa-Sakurai, Mari Murai-Hatano, Arifa Ahamed, and Matsuo Uemura. Aquaporins in developing rice grains. Bioscience, Biotechnology, and Biochemistry, 79(9):1422–1429, 2015.

[62] Noriko Kinoshita, Masayuki Kato, Kei Koyasaki, Takuya Kawashima, Tsutomu Nishimura, Yuji Hirayama, Itsuro Takamure, Takashi Sato, and Kiyoaki Kato. Identification of quantitative trait loci for rice grain quality and yield-related traits in two closely related oryza sativa l. subsp. japonica cultivars grown near the northernmost limit for rice paddy cultivation. Breeding Science, page 16155, 2017.

[63] Xiaoqiong Li, Yu Wei, Jun Li, Fangwen Yang, Ying Chen, Yinhua Chen, Sibin Guo, and Aihua Sha. Identification of qtl tgw12 responsible for grain weight in rice based on recombinant inbred line population crossed by wild rice (oryza minuta) introgression line k1561 and indica rice g1025. BMC Genetics, 21(1):10, 2020.

[64] S Sreedhar, T Dayakar Reddy, MS Ramesha, et al. Genotype x environment interaction and stability for yield and its components in hybrid rice cultivars (oryza sativa l.). International Journal of Plant Breeding and Genetics, 5(3):194–208, 2011.

[65] KA Steele, Adam Huw Price, HE Shashidhar, and JR Witcombe. Marker-assisted selection to introgress rice qtls controlling root traits into an indian upland rice variety. Theoretical and Applied Genetics, 112(2):208–221, 2006.

[66] KK Jena and DJ Mackill. Molecular markers and their use in marker-assisted selection in rice. Crop Science, 48(4):1266–1276, 2008.

[67] Zhe Zhang, Ulrike Ober, Malena Erbe, Hao Zhang, Ning Gao, Jinlong He, Jiaqi Li, and Henner Simianer. Improving the accuracy of whole genome prediction for complex traits using the results of genome wide association studies. PloS One, 9(3), 2014.

[68] Qifa Zhang, Jiayang Li, Yongbiao Xue, Bin Han, and Xing Wang Deng. Rice 2020: a call for an international coordinated effort in rice functional genomics. Molecular Plant, 1(5):715–719, 2008.

[69] ANM Rubaiyath Bin Rahman and Jianhua Zhang. Rayada specialty: the forgotten resource of elite features of rice. Rice, 6(1):41, 2013.

[70] Antonio T Perez and Muhammad Nasiruddin. Field notes on the rayadas: a flood-tolerant deepwater rice of bangladesh. In Proceedings of the International Seminar on Deepwater Rice, number 15, pages 87–91, 1974.

[71] Jean-Christophe Glaszmann. Isozymes and classification of asian rice varieties. Theoretical and Applied Genetics, 74(1):21–30, 1987.

